# Arginine shortage induces replication stress and confers genotoxic resistance by inhibiting histone H4 translation and promoting PCNA polyubiquitination

**DOI:** 10.1101/2023.01.31.526362

**Authors:** Yi-Chang Wang, Andrew A. Kelso, Adak Karamafrooz, Yi-Hsuan Chen, Wei-Kai Chen, Chun-Ting Cheng, Yue Qi, Long Gu, Linda Malkas, Hsing-Jien Kung, George-Lucian Moldovan, Alberto Ciccia, Jeremy M. Stark, David K Ann

## Abstract

The unique arginine dependencies of cancer cell proliferation and survival creates metabolic vulnerability. Here, we investigate the impact of extracellular arginine availability on DNA replication and genotoxic resistance. Using DNA combing assays, we find that when extracellular arginine is limited, cancer cells are arrested at S-phase and DNA replication forks slow or stall instantly until arginine is re-supplied. The translation of new histone H4 is arginine-dependent and impacts DNA replication and the expression of newly synthesized histone H4 is reduced in the avascular nutrient-poor breast cancer xenograft tumor cores. Furthermore, we demonstrate that increased PCNA occupancy and HLTF-catalyzed PCNA K63-linked polyubiquitination protects arginine-starved cells from hydroxyurea-induced, DNA2-catalyzed nascent strand degradation. Finally, arginine-deprived cancer cells are tolerant to genotoxic insults in a PCNA K63-linked polyubiquitination-dependent manner. Together, these findings reveal that extracellular arginine is the “linchpin” for nutrient-regulated DNA replication. Such information could be leveraged to expand current modalities or design new drug targets against cancer.

## INTRODUCTION

A major hallmark of tumor progression is unchecked DNA replication and adaptation to the high metabolic demands of rapid cell division in a nutritionally deprived microenvironment (Boroughs and DeBerardinis, 2015; Lama-Sherpa and Shevde, 2020). Arginine, a semi-essential amino acid, is a key nutrient (Chen et al., 2021a) for arginine auxotrophic tumors (Cheng et al., 2018; Lee et al., 2018). These tumors become reliant on external sources of arginine due to widespread urea cycle dysregulation (Lee et al., 2019; Pan et al., 2016; Wu, 2009). External arginine shortages in the tumor core can result from poor vascularization and dysfunctional blood flow in the tumor microenvironment (TME) (Lee et al., 2019; Pan et al., 2016). Furthermore, limiting dietary arginine or use of arginine-deprivation agents leads to the inhibition of arginine auxotroph tumor growth (Chen et al., 2021b; Cheng et al., 2018; Hsu et al., 2021; Poillet-Perez et al., 2018; Qiu et al., 2014). Moreover, the spatiotemporal regulation of DNA replication during S-phase of the cell cycle, is synchronized with nutrient availability to cells (Bohnsack and Hirschi, 2004; Cuyas et al., 2014; Liu and Sabatini, 2020). However, whether extracellular arginine shortage represents a widespread obstacle to DNA replication and how DNA replication contends with insufficient extracellular arginine availability remains largely unknown.

During DNA replication, nucleosomes are disrupted as the replication forks pass, followed by reassembly of recycled parental histones and incorporation of newly synthesized histones to restore chromatin state (MacAlpine and Almouzni, 2013). As such, the synthesis of replication-dependent histones, essential building blocks for chromatin reassembly behind the replication fork, is tightly coupled to ongoing DNA synthesis (Hammond et al., 2017). Indeed, histone biosynthesis is S-phase-specific to govern S-phase timing and cell cycle progression (Armstrong and Spencer, 2021; Gunesdogan et al., 2014; Mejlvang et al., 2014). Given that arginine is a proteinogenic amino acid, it is clearly important to understand the role of arginine depletion on replication-dependent histone biosynthesis.

DNA replication stress comprises a multitude of cellular conditions, including physical blockage of DNA replication fork progression along the template, deregulation of the replication initiation or elongation complexes, or deoxyribonucleotide depletion (Zeman and Cimprich, 2014). Proliferating cell nuclear antigen (PCNA) plays a vital role in coordinating DNA replication activity and maintaining genome integrity (Berti et al., 2020; Moldovan et al., 2007). PCNA serves as a hub for proteins involved in DNA synthesis, cell cycle control, and DNA damage response and repair (Kanao and Masutani, 2017; Kelman, 1997; Komatsu et al., 2000; Ulrich, 2009). PCNA is also subjected to multiple posttranslational modifications, including ubiquitination, contributing to the coordination of DNA replication and genotoxic damage tolerance (Papouli et al., 2005; Ulrich and Takahashi, 2013). Whereas PCNA monoubiquitination in response to DNA damage promotes the bypass of replication-blocking lesions (Kannouche et al., 2004), PCNA polyubiquitination has been associated with fork reversal (Hoege et al., 2002). In particular, the mammalian RAD5 ortholog helicase-like transcription factor (HLTF), an ubiquitin ligase for PCNA, has also been shown to be required for replication fork reversal (Bai et al., 2020; Kile et al., 2015). However, mechanisms by which arginine levels affect PCNA levels on replicating chromatin and the factors involved in this process remain elusive. In addition, it is currently unknown how arginine availability impacts DNA replication stress response and adaptation to genotoxic insult.

Here, we report that extracellular arginine shortage suppresses DNA replication elongation by reducing newly synthesized histone H4. Transient transfection of recombinant histone H4 protein can partially rescue DNA replication in arginine-starved cells. We present evidence that both HLTF and PCNA play key roles in fork protection in the absence of arginine. Arginine shortage increases HLTF-mediated PCNA K63-linked polyubiquitination and PCNA occupancy on the stalled nascent DNA strands. Next, we find that under replication stress induced by hydroxyurea (HU), arginine-starved cells exhibit vigorous nascent strand degradation, when HLTF is depleted, in a manner dependent on DNA2. Finally, HLTF depletion or treatment with PCNA ligand AOH996 results in a higher DNA damage signaling and sensitivity to genotoxicity under arginine shortage. Our results highlight a previously undescribed replication stress that arginine shortage induces defects on *de novo* chromatin assembly to interfere with DNA replication and S-phase progression. On the basis of these results, we conclude that extracellular arginine shortage triggers replication stress responses and confers tumor escape from a variety of genotoxicities.

## RESULTS

### Extracellular arginine regulates DNA replication fork progression

To determine the fate of cells that experience extracellular arginine shortage during cell cycle progression, asynchronously cycling arginine auxotrophic MDA-MB-231 breast cancer cells were pulse-labeled with bromodeoxyuridine (BrdU; t = 0 h), an analog of thymidine, and then chased in medium ± arginine for an additional 10 h. Using flow cytometry, we found that in the presence of arginine, most BrdU-labeled MDA-MB-231 cells were not at S-phase (**Fig. 1A**, *lower middle panel*) while cells in arginine-free media were retained at S-phase (**Fig. 1A**, *lower right panel*). Multiple cell lines, irrespective of arginine auxotrophy, displayed a similar impediment to S-phase exit in response to extracellular arginine shortage (**Fig. 1A** *right panel*). The overall cell cycle distribution of G1, S and G2/M were analyzed, and increased S-phase and decreased M-phase populations were found in arginine-deprived cells (**Fig. S1A**). To determine if this stalled S-phase phenomenon occurs within the S-phase, we synchronized MDA-MB-231 cells in early S-phase using a double thymidine block. BrdU-labeled synchronized cells did not exit S-phase when deprived of arginine (**Fig. 1B**). Further, hallmarks of cell cycle progression such as the upregulation of Cyclin A, Cyclin B1, phosphorylated H3S10 and H4 were absent in cells released in arginine-free medium, further confirming the inhibitory effect of arginine shortage on S-phase progression (**Fig. S1B**). This observed S-phase blockade induced by arginine removal was reversible when 2.6 to 5.3 mg/L arginine was added to the medium (**Fig. S1C**). In parallel, the overall incorporation of 5-ethynyl-2’-deoxyuridine (EdU) analog in breast cancer-derived BT-549 spheroids decreased in the absence of arginine in a time-dependent manner (**Fig. S1D**). We conclude that arginine is required for the cell cycle progression of S-phase cells.

**Fig. 1.**
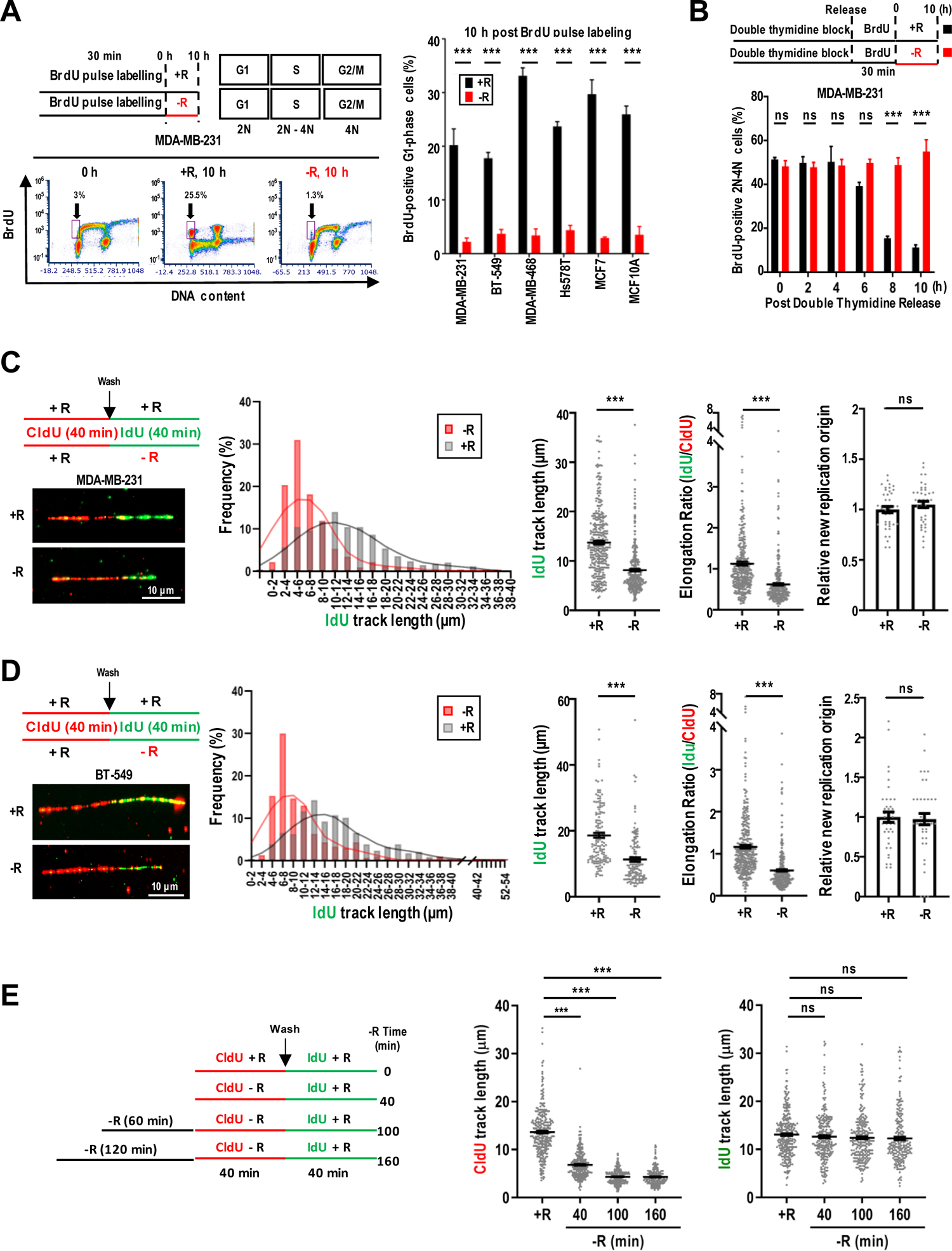
Arginine is indispensable for DNA replication. (**A**) Cell cycle progression analysis after BrdU labeling under ± arginine conditions using flow cytometry. *Top left*: Experimental outline. *Bottom left*: Analysis of BrdU^+^ MDA-MB-231 cells during G1-phase (purple-boxed). *Right:* Quantification of percentages of BrdU^+^ G1 cells in multiple cancer cell lines. N = 3 independent experiments. At least 50,000 events were collected and analyzed. (**B**) S-phase analysis in double thymidine-synchronized MDA-MB-231 cells pulse-labeled with BrdU under ± arginine conditions using flow cytometry. *Top panel*: Experimental outline. *Bottom*: Quantification of percentages of BrdU^+^ S-phase cells at different time periods after thymidine block release. N = 3 independent experiments. At least 50,000 events were collected and analyzed. (**C**, **D**) DNA fiber analysis using CldU (red) or IdU (green) antibodies under ± arginine conditions in MDA-MB-231 (**C**) or BT-549 (**D**) cells. *Top left*: Experimental design. *Bottom left*: Representative images of DNA fibers. *Middle:* Histograms shows percentage frequency of IdU track length. *Right*: Quantitation of IdU track length, elongation ratio (IdU/CldU) and firing of relative new replications of origin. Each dot represents one fiber or one field. At least 100 tracts or 40 fields were analyzed per sample. (**E**) DNA combing analysis in MDA-MB-231 cells deprived of arginine for varying time-frames and then re-supplemented with arginine (84 mg/L, the same amount of arginine in full medium). *Left*: Experimental outline. *Right*: Quantitation of CldU and IdU track lengths. N = number of fields. At least 100 tracts were analyzed per sample. Mean ± SEM is shown; ns: not significant; ***: *p*<0.005 (two-tailed unpaired Student’s *t*-test); R: arginine.

We hypothesized that extracellular arginine is required for DNA replication fork progression. To address this possibility, we studied replication fork dynamics in cells at single-molecule resolution under ± arginine conditions. First, MDA-MB-231 and BT-549 cells were pulse-labeled with thymidine analog, 5-chloro-2’-deoxyuridine (CldU), for 40 min to establish the baseline tract lengths. After washing, cells were concomitantly labeled with the second thymidine analog iododeoxyuridine (IdU) under ± arginine conditions for 40 min (**Fig. 1C, 1D**, *upper left diagrams*). The track length of ongoing replication forks (with adjacent CldU (1^st^ labeling, red) and IdU (2^nd^ labeling, green) signals) was examined and quantified (**Fig. 1C, 1D**, *lower left panels*). Quantitative analyses confirmed that the frequency of IdU-labeled tracts with reduced lengths increased in both cell lines upon arginine shortage, but this was not due to the new initiation of DNA replication (**Fig. 1C, 1D**, *right 4 panels*).

Next, we determined whether the duration of arginine shortage affects DNA replication fork resumption after arginine is re-supplied. Cells that were deprived of arginine for up to 120 min were first pulse-labeled with CldU in arginine-free medium for an additional 40 min (i.e., up to 160 min), and subsequently pulse-labeled with IdU in arginine-containing medium for 40 min (**Fig. 1E**, *left diagram*). Notably, we found a time-dependent shortening of CldU-labeled tracts up to 100 min of arginine shortage (**Fig. 1E**, *middle panel*). However, arginine replenishment restored DNA fork progression, visualized with IdU-labeled tracks and track length, despite varying arginine-free exposure times (**Fig. 1E**, *right panel*). This indicates that transient extracellular arginine shortage is not a catastrophic DNA replication stress event, and replication forks are primed to resume upon arginine re-addition. Together, we propose that arginine shortage holds DNA replication forks in a “standby” mode until the arginine shortage is resolved, and concomitantly arginine shortage decelerates DNA elongation in an immediate, yet reversible, manner.

### Extracellular arginine controls DNA replication by promoting the synthesis of histone H4 protein

Since arginine is a proteogenic amino acid and the biosynthesis of core histone proteins are tightly coupled to the cell cycle and peaks specifically during S-phase, concomitant with DNA replication (Gunesdogan et al., 2014; Marino-Ramirez et al., 2006), we surveyed whether arginine promotes DNA replication by regulating translation. We adopted the AHA (L-azidohomoalaine, a methionine analog) incorporation assay to assess the direct impact of arginine shortage on newly synthesized proteins in S-phase cells (**Fig. S2A**). While those Coomassie blue stained signals were only subtly affected when cells were released in arginine-free medium (**Fig. S2B,** *left 2 panels*), AHA-labeled proteins were markedly reduced upon arginine shortage (**Fig. S2B,** *right 2 panels*). To determine the effect of arginine availability on newly synthesized histone H2B, H3 and H4, AHA-labeled proteins were isolated using a biotin pull-down strategy followed by Western analyses. As shown in **Fig. 2A**, histone H4, unlike H2B and H3, was not synthesized upon arginine shortage. In stark contrast, AHA-labeled ATF4 was induced, serving as an arginine shortage-induced ER stress control (Cheng et al., 2018). To rule out the possibility that arginine shortage also reduced histone H4 mRNA abundance, qRT-PCR analysis was performed. We found that H4 mRNA abundance began to rise at 30 min after released from double thymidine block, irrespective of arginine availability, validating that cells are in S-phase (**Fig, S2C**). However, H4 mRNA level subtly leveled off between 4 h and 8 h post-release in full medium, rendering it significantly lower than H4 mRNA in arginine-deprived cells (**Fig, S2C**). These results suggest that arginine shortage leads to a preferential depletion of newly synthesized histone H4 via a translational control.

**Fig. 2.**
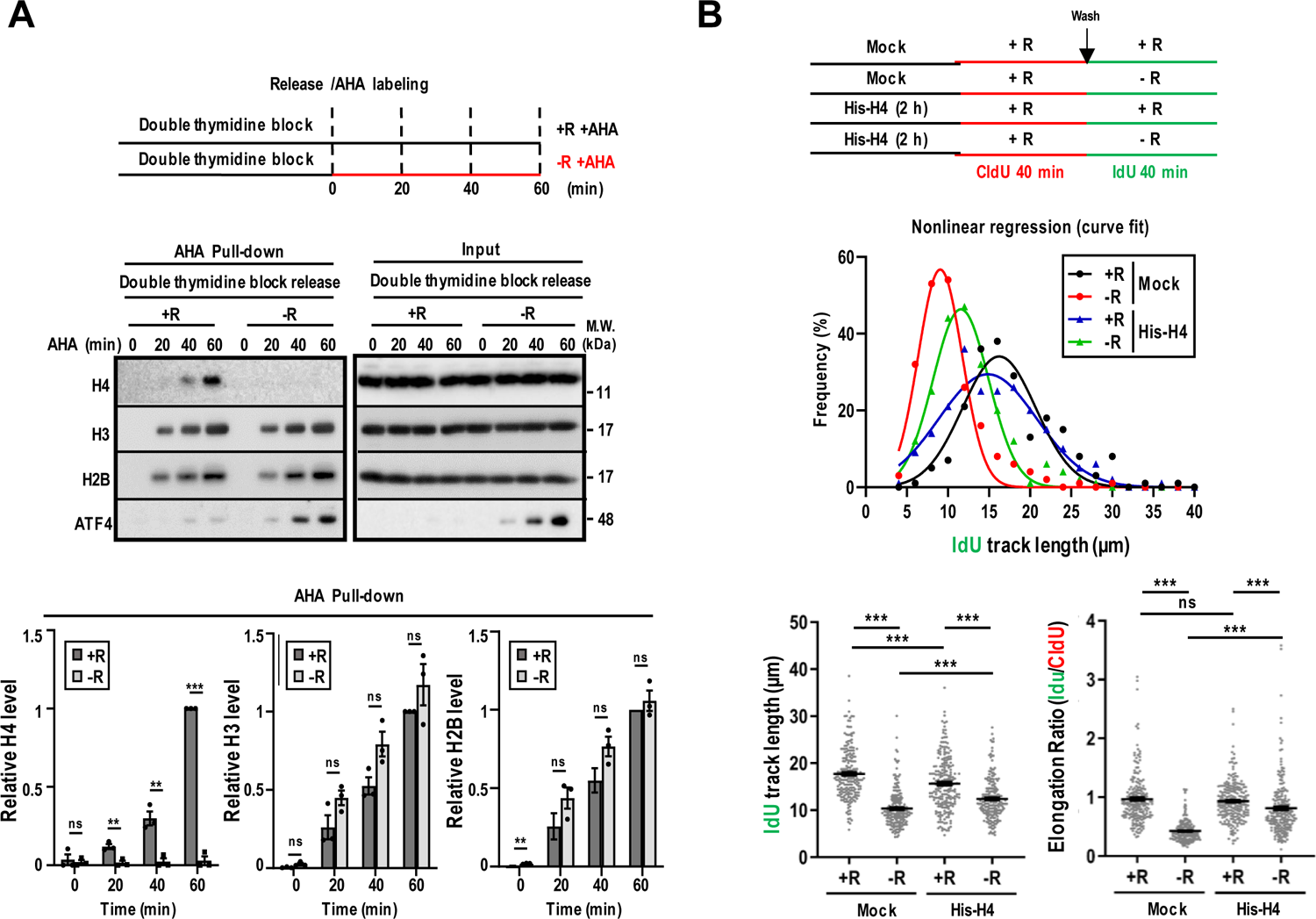
Transfected recombinant histone H4 protein rescues DNA elongation under arginine shortage. (**A**) Analysis of nascent AHA-labeled histones in S-phase enriched MDA-MB-231 cells. *Top*: Experimental outline. *Middle*: Representative Western blot*. Bottom:* Densitometric quantification of relative levels of histones and ATF4. Relative protein levels were calculated by dividing the intensity of desired band with that in cells released to +R medium for 60 min (set as 1). N = 3 independent experiments were analyzed. (**B**) DNA combing assay using MDA-MB-231 cells transfected with His-tagged histone H4 under ± arginine conditions. *Top:* Experimental outline. *Middle:* Nonlinear regression curve analysis for IdU track length. *Bottom:* Quantification of IdU track lengths and IdU/CldU ratio. At least 200 tracts were analyzed per sample. Mean ± SEM is shown; ns: not significant; ***: *p*<0.005 (two-tailed unpaired Student’s *t*-test).

Next, we tested whether increased newly synthesized H4 rescues DNA replication in arginine-deprived cells. To this end, we used a protein transfection reagent to introduce recombinant human His-tagged histone H4 proteins into cells, modeling an increase in newly synthesized H4, prior to arginine removal (**Fig. S2D**, *left panel*). After confirming the incorporation of recombinant histone H4 protein into chromatin (**Fig. S2D**, *right panel*), DNA combing assays showed that transfected recombinant H4 protein, at least in part, rescued both IdU-labeled track lengths and IdU/CldU ratio in arginine-depleted cells (**Fig. 2B**). Together, our data suggest that while histone H4 is not the only protein involved in DNA replication whose synthesis is selectively suppressed by arginine depletion, the transfected recombinant histone H4 is able to partially phenocopy extracellular arginine to promote DNA replication. This also explains why extracellular arginine shortage prevents S-phase exit (**Fig. 1A, 1B**).

### Newly synthesized histone H4 marks are surrogate markers of arginine availability within the TME

To search for a marker for arginine availability and/or DNA replication speed, we exposed cells to various arginine concentrations for different durations and assessed histone H4 marks via Western blot analysis. Both H4K5ac and H4K12ac marks, which are enriched at newly assembled chromatin (Armstrong and Spencer, 2021), decreased in MDA-MB-231 (**Fig. 3A**) and BT-549 cells (**Fig. S3A**) when exposed to suboptimal extracellular levels of arginine. Multiple cell lines displayed similar results upon 2 h of extracellular arginine shortage, regardless of their *de novo* arginine biosynthesis capacity via ASS1 (Keshet et al., 2018) (**Fig. S3B**). Because newly translated histone H4 is acetylated on lysine 5 and 12 in the cytoplasm prior to nuclear translocation, accumulation and nucleosome deposition (Denu, 2019; Sobel et al., 1995; Taddei et al., 1999), we assessed the steady-state levels of H4K5ac and H4K12ac in the cytoplasmic and nuclear subfractions of cells released into ± arginine medium for 4 h post-double thymidine block. The lack of nuclear lamin A/C and cell surface CD44 signals in the cytoplasmic and nuclear fractions, respectively, confirmed the relative purity of each fraction. Following arginine shortage, the overall levels of lamin A/C, CD44, histone H3 and actin remained unchanged, and the nuclear accumulation of ATF4 served as an arginine shortage control (**Fig. 3B, S3C**) (Cheng et al., 2018). Consistently, a universal decrease of newly synthesized histone H4 marks in total cell lysates, cytoplasmic and nuclear fractions was observed with arginine shortage (**Fig. 3B, S3C**). This arginine removal-mediated decline of histone marks was more notable in H4K5ac and H4K12ac (∼50%), but not in H4K16ac, H3K9me3, and H3K4me3 levels (**Fig. S3D**). To support that the arginine shortage contributes to the reduction of newly synthesized histone H4 marks, supplementation of arginine rapidly restored H4K5ac and H4K12ac signals (**Fig. S3E**), correlating well with DNA replication resumption (**Fig. 1E**).

**Fig. 3.**
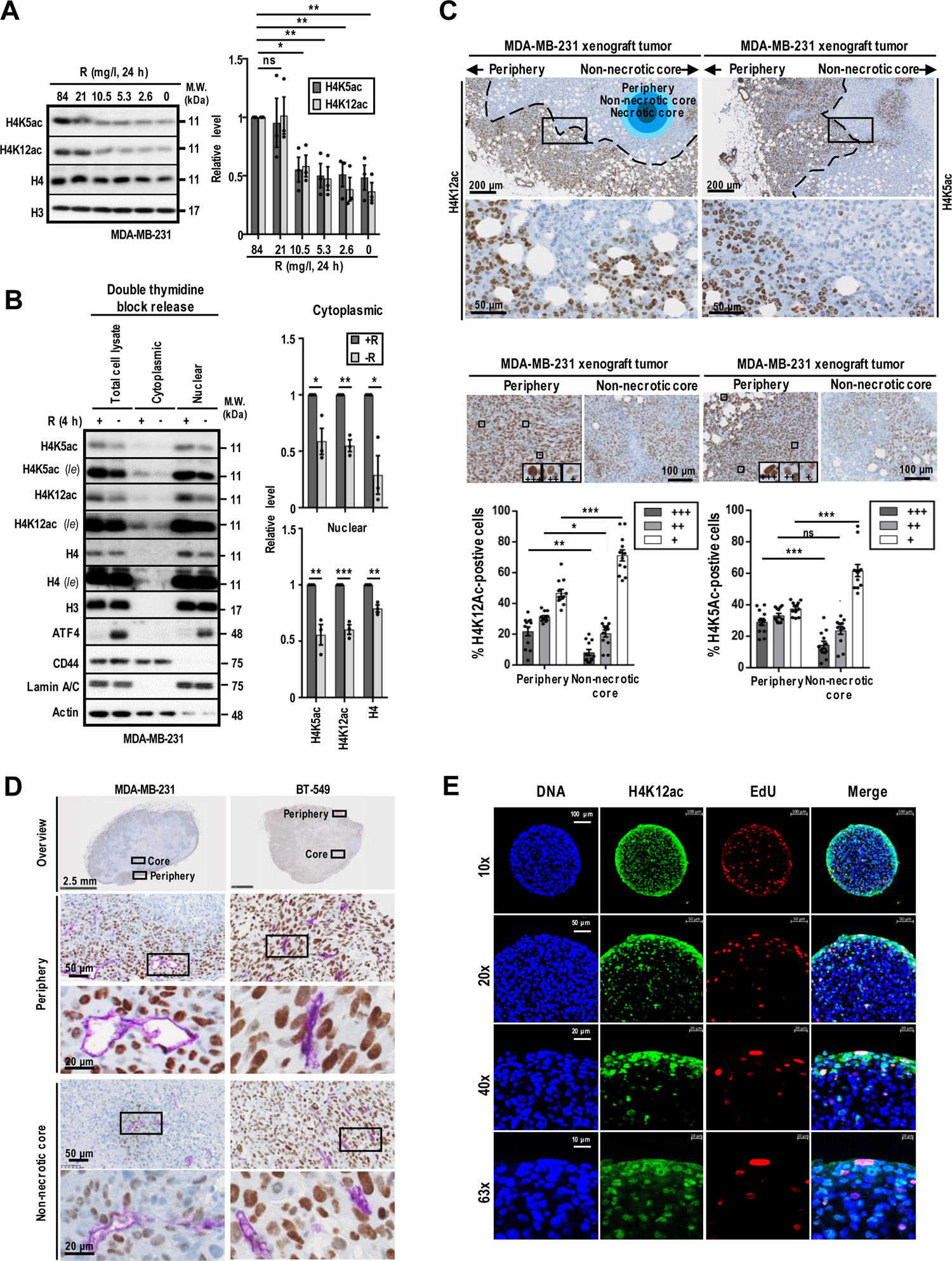
Newly synthesized histone H4 marks are highly correlated with arginine availability. (**A**) Analysis of H4K5ac and H4K12ac in MDA-MB-231 cells grown under varying arginine conditions. *Left:* Representative Western blot. *Right:* Densitometric quantification of relative levels of H4K5ac and H4K12ac normalized to control (84 mg/l). N = 3 independent experiments. (**B**) Subcellular fraction analysis of H4K5ac and H4K12ac levels in synchronized S-phase MDA-MB-231 cells grown under ± arginine conditions. *Left:* Representative Western blot. *Right:* Densitometric quantification of relative levels of H4K5ac and H4K12ac normalized to control (+R). N = 3 independent experiments. (**C**) H4K12ac and H4K5ac expression in MDA-MB-231 xenograft tumors. *Top:* Representative IHC image of periphery and non-necrotic core and corresponding enlarged view of inset area (black square) shows staining for H4K12ac and H4K5ac (brown). Dashed line: Boundary between periphery and non-necrotic core regions. *Bottom:* Quantification of the intensities of H4K12ac^+^ and H4K5ac^+^ MDA-MB-231 cells from representative IHC images (N = 12) and the quantification of the respective percentage of cells with indicated H4K5ac or H4K12ac intensity. Insets in images show +++ strong, ++ medium, and + weak positive staining. (**D**) Representative IHC images of CD31 (purple) and H4K12ac (brown) expression in MDA-MB-231 and BT-549 xenograft tumors. An overview of full tumor section is shown (*top*). A corresponding enlarged view of boxed area is shown below for the periphery and the non-necrotic core. (**E**) Representative immunofluorescence image of a BT-549 tumor spheroid cryosection stained for H4K12ac (green), EdU (red) and Hoechst 33342 (Blue). experiments. Mean ± SEM is shown; ns: not significant; *: *p*<0.05; **: *p*<0.01; ***: *p*<0.005; two-tailed unpaired Student’s *t*-test; *le*: long exposure; R: arginine.

A typical feature of solid tumors is their heterogeneous distribution of blood vessels and flow (Rausch et al., 2017). Consequently, suboptimal nutrient availability develops within the avascularized TME. Previous studies have shown that arginine is one of the least available amino acids in the tumor core regions (Pan et al., 2016). To determine how DNA replication is perturbed by the lack of arginine supply *in vivo*, we sought to determine whether the distribution of H4K5ac and H4K12ac marks are located together with cell proliferation marker Ki67 in breast cancer MDA-MB-231 and BT-549 xenograft tumors using immunohistochemical (IHC) staining. Quantitative analyses confirmed, in general, the gradual decrease of H4K5ac and H4K12ac signals within the non-necrotic tumor core compared to the peripheral regions of MDA-MB-231 (**Fig. 3C**) and BT-549 (**Fig. S4A**) xenograft tumors. In addition, CD31-positive cells, an endothelial cell marker, and H4K12ac-positive cells were detected in close proximity at the periphery and non-necrotic core regions (**Fig. 3D**), and H4K12ac-negative cells were largely enriched in the CD31-negative regions (**Fig. S4B**). In addition, S-phase marker Ki67 was also enriched at the peripheral region, correlating well with H4K12ac (**Fig. S4C**) in MDA-MB-231 and BT-549 xenografts. Consistent with these observations, most EdU-labeled cells were localized to the periphery of BT-549 spheroids (**Fig. 3E**). Likewise, most H4K12ac-positive cells were found at the peripheral of Hs578T and MCF-7 tumor spheroids (**Fig. S4D**). To test the uniqueness of arginine shortage, we deprived other nutrients, such as glutamine, which is limited in tumor core regions, and serum as well as leucine, which are indispensable for cell proliferation. Results showed that neither removal of serum, glutamine or leucine affected H4K5ac and H4K12ac signals (**Fig. S4E-G**) (Liu et al., 2014; Pan et al., 2016). Collectively, these data suggest that extracellular arginine availability preferentially affects H4K5ac and H4K12ac abundance *in vitro* and *in vivo*.

### Extracellular arginine shortage creates genotoxic tolerance

The maintenance of genome stability relies on successful DNA replication and proper DNA damage response in S-phase. Next, we hypothesize that arginine availability might impact the DNA replication stress response and/or cell recovery from genotoxicity. When replication fork stress or DNA damage occurs, single-stranded DNA (ssDNA) is created and elicits the DNA damage response (Cimprich and Cortez, 2008; Zeman and Cimprich, 2014). RPA-coating of ssDNA results in the activation of the ATR/Chk1 pathway to promote replication fork stability and genome stability (Cimprich and Cortez, 2008; Zeman and Cimprich, 2014). To test our hypothesis, we monitored the replication stress response via the phosphorylation of RPA (p-RPA(S33)) and Chk1 (p-Chk1(S345)) in arginine-starved cells. We used hydroxyurea (HU) and camptothecin (CPT) as positive controls, both are known agents that cause DNA replication stress and activates the ATR/Chk1 pathway (**Fig. 4A**) (Cimprich and Cortez, 2008; Zeman and Cimprich, 2014). Arginine shortage (2 h) failed to induce p-Chk1(S345) and phosphorylation of RPA in multiple cell lines (**Fig. 4A, S5A**). Consistent with this, we failed to observe an increase in the chromatin-bound RPA levels, a reflection of the degree of ssDNA exposure, in pre-extracted S-phase cells via flow cytometry post-arginine shortage (4 h) (**Fig. 4B**). This is in contrast with the observations following HU- and CPT-treatment (Forment and Jackson, 2015). The inability of extracellular arginine shortage to activate Chk1 phosphorylation and RPA chromatin binding suggests that arginine shortage did not lead to significant ssDNA accumulation while deaccelerating DNA replication.

**Fig. 4.**
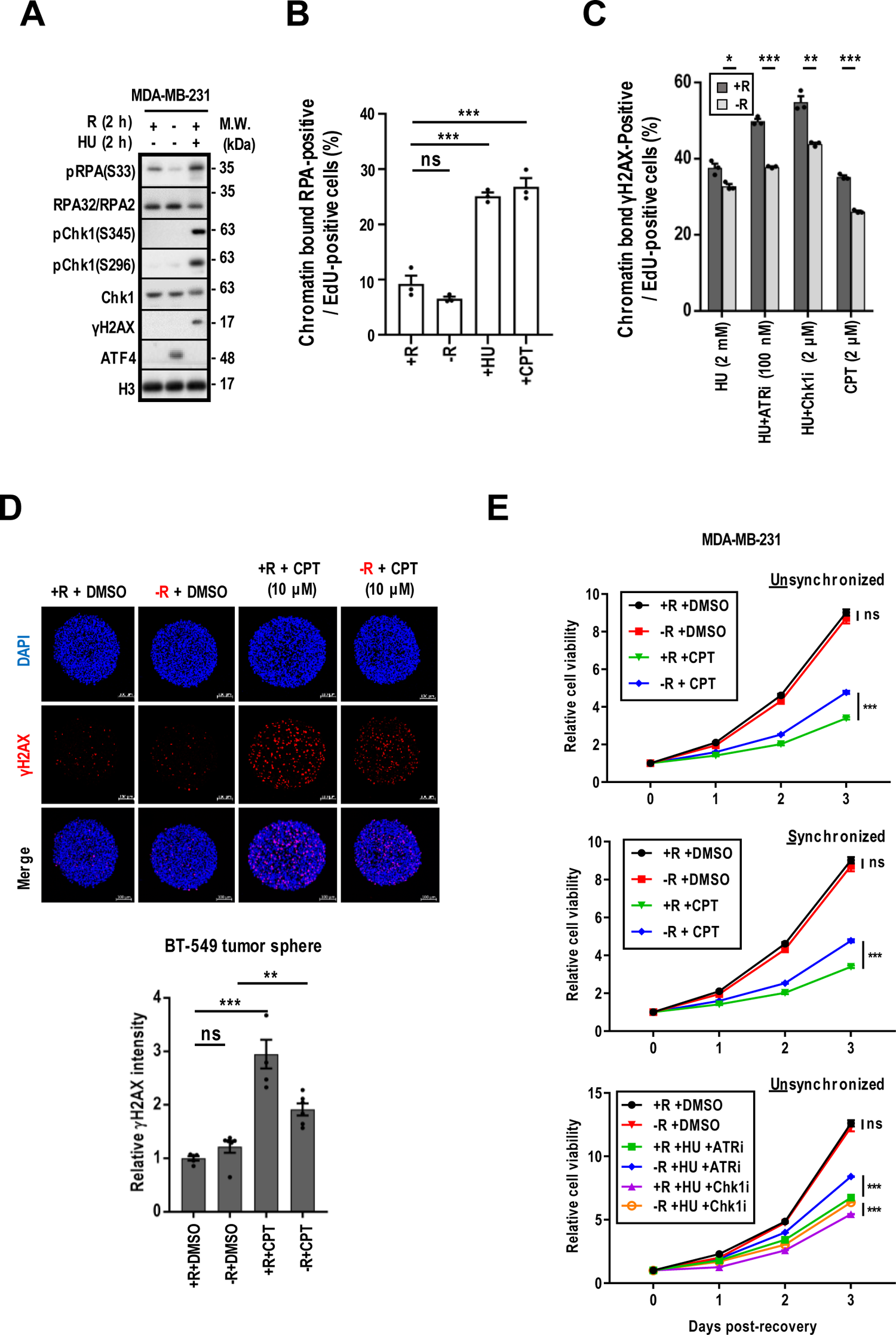
Arginine shortage promotes resistance to genotoxic insult. (**A**) Representative Western blot for RPA (S33) and Chk1 (S345) phosphorylation in HU-treated MDA-MB-231 cells under ± arginine conditions. N = 3 independent experiments. (**B, C**) Quantification of chromatin-bound RPA^+^ (**B**) and γH2AX^+^ (**C**) EdU-labeled MDA-MB-231 cells grown under stated conditions for 4 h following flow cytometry analysis. N = 3 independent experiments. At least 10,000 events were collected and analyzed. (**D**) γH2AX expression analysis in BT-549 spheroids treated under stated conditions for 4 h. *Top:* Representative fluorescent images of BT-549 spheroid cryosections stained with DAPI (blue) and γH2AX (red). *Bottom:* Quantification of relative γH2AX intensity normalized to tumor area. N = 4 independent experiments. (**E**) Analysis of relative cell recovery of MDA-MB-231 cells under stated conditions for 4 h followed by recovering in regular full medium. Cell viability was analyzed at indicated time points. Mean ± SEM is shown; two-tailed unpaired Student’s *t*-test; ns: not significant; *: *p*<0.05; **: *p*<0.01; ***: *p*<0.005; CPT: camptothecin; HU: hydroxyurea; R: arginine.

We next determined whether the DNA damage response to double-strand DNA breaks (DSBs) was impacted under ± arginine conditions by assessing the histone variant H2AX phosphorylation on S139 (γH2AX) (Sirbu et al., 2011) in MDA-MB-231 cells. To induce DSBs we used: (1) CPT alone (Ramaswamy et al., 1987), and (2) HU, a recognized source of DNA damage by misincorporation (Techer et al., 2017), in combination with either Chk1 inhibitor (Chk1i, rabusertib) (van Harten et al., 2019)) or an ATR inhibitor (ATRi, BAY1895344) (Kawasumi et al., 2014). Chk1i (1 μM) alone, but not ATRi (100 nM) alone, induced p-Chk1(S345) without a marked γH2AX accumulation (**Fig. S5B**, *lane #7 vs. #5*), and co-treatment of ATRi (100 nM) blocked HU-induced p-Chk1(S345), but induced γH2AX (**Fig. S5B**, *lane #9 vs. #3*), serving as a positive control (Leung-Pineda et al., 2006). As expected, arginine depletion reduced γH2AX chromatin binding in S-phase cells, treated with genotoxic agents, analyzed by flow cytometry (**Fig. 4C**). Notably, arginine shortage attenuated ATRi- or Chk1i-sensitized γH2AX accumulation in HU-treated MDA-MB-231 cells (**Fig. S5B**, *lane #9 vs. #10 and lane #11 vs. #12*) and γH2AX signals in tumor spheroids treated with CPT (**Fig. 4D**). Taken together, cells exposed to arginine shortage appeared to exhibit reduced DNA replication stress responses than cells maintained in full medium upon treatment with the same genotoxic agents.

To determine the impact of arginine on cell recovery from genotoxic treatments, MDA-MB-231 cells were subjected to CPT (2 μM), HU (2 mM)+ATRi (100 μM) or HU (2 mM)+Chk1i (1 μM) under ± arginine conditions for 4 h, and then cultured in drug-free full medium for indicated time periods, with or without S-phase synchronization (**Fig. 4E**). Similar results were found in CPT-treated BT-549 and MCF-7 cells (**Fig. S5C**). There was a significant gain in resistance to genotoxic agents tested in arginine-starved cells at day 3 post-treatment. A similar result was observed in tumor spheroids (**Fig. S5D**). Collectively, unlike other DNA replication stalling chemicals which induce DNA damage responses, arginine shortage slows down DNA replication without causing DNA damage responses while enabling tolerance to genotoxic insults.

### Distinct protein dynamics at arginine shortage-stressed replication forks

We next sought to identify arginine shortage-induced and ATR-independent signaling events that mediate DNA replication slowing in MDA-MB-231 cells using the accelerated isolation of proteins on nascent DNA (aniPOND) method (Leung et al., 2013). Due to the generation of shorter EdU-incorporated nascent DNA strands under arginine removal (**Fig. 1C, D**), we expected less H4K5ac, H4K12ac, H2B, H3 and H4 to be pulled down from EdU-labeled cells exposed to arginine shortage compared to cells grown in full medium (**Fig. 5A**). Furthermore, arginine replenishment following shortage restored these histone levels to similar levels observed in cells grown in full medium (**Fig. 5A**). Decreased levels of H4K5ac and H4K12ac marks showed that there was less newly synthesized H4 occupying nascent DNA strands during DNA replication under arginine shortage (**Fig. 5A**). In stark contrast with reduced histone levels, we found that more PCNA was bound to EdU-labeled DNA after arginine removal (**Fig. 5B**) and this did not occur with arginine re-supply following arginine shortage (**Fig. 5B**). As previously reported (Betous et al., 2018; Sirbu et al., 2011), CPT- and HU-treatment did not increase nascent DNA-bound PCNA levels (**Fig. 5B**), serving as controls. Furthermore, chromatin fractionation experiments revealed that a higher level of chromatin-bound PCNA was exclusively detected in arginine-deprived cells, and both CPT- and HU-treatments failed to affect occupancy of chromatin-bound PCNA under ± arginine conditions (**Fig. S6A**). These results suggested that there was increased occupancy of PCNA on nascent DNA strands at stalled replication forks upon arginine withdrawal.

**Fig. 5.**
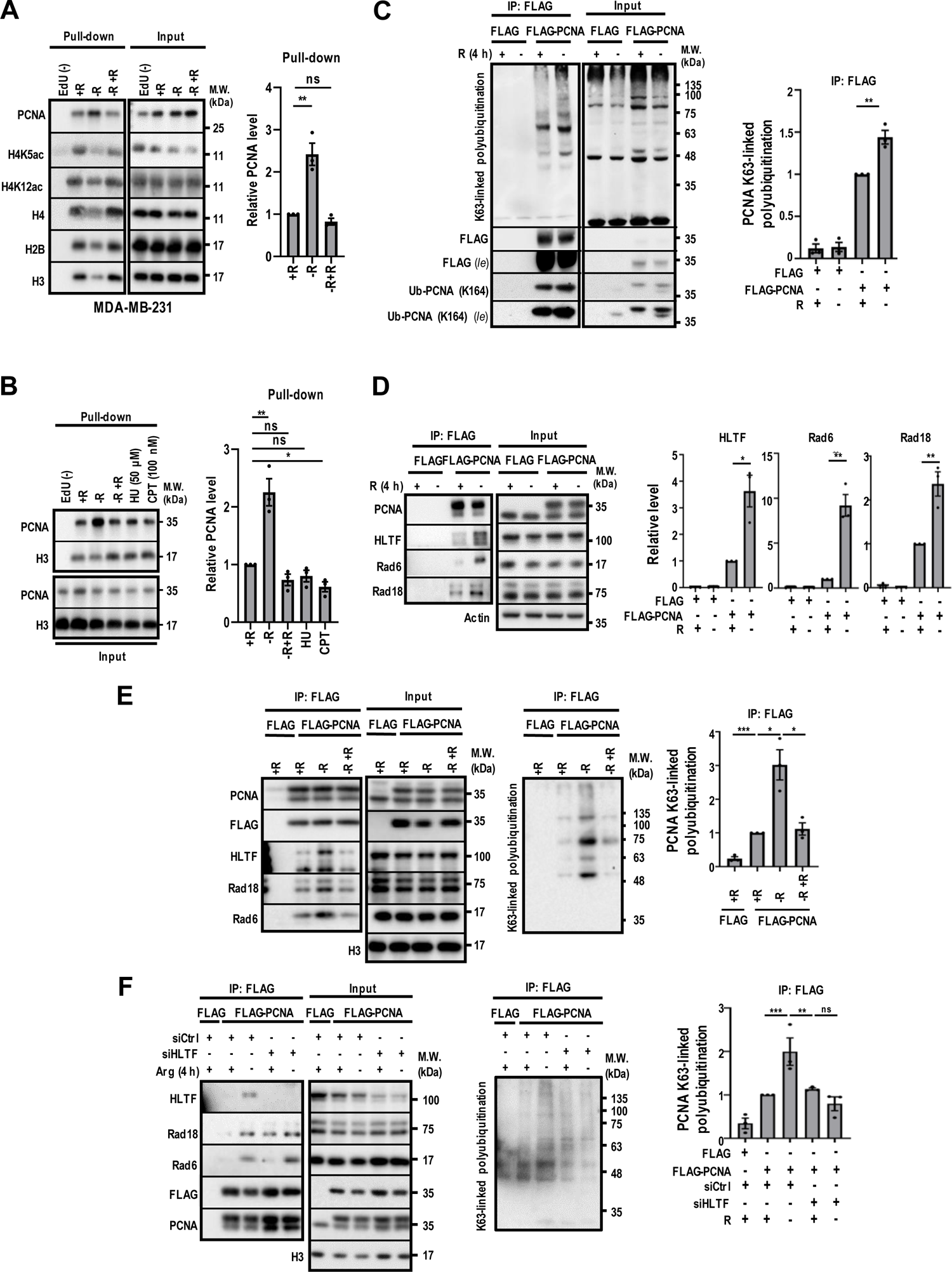
Arginine shortage-dependent regulation of PCNA and HLTF. (**A-B**) Representative Western blots of fork-associated proteins isolated using aniPOND method from EdU^+^ MDA-MB-231 cells grown under stated conditions. EdU (-): negative control. N = 3 independent experiments. (**C-F**) Western blot analysis of immunoprecipitated PCNA-associated proteins from Myc-ubiquitin transfected FLAG-PCNA-overexpressed HEK293T cells and subjected to indicated treatments. *Left*: Representative Western blots are shown for the indicated antibodies. *Right*: Densitometric quantification of relative levels of indicated proteins normalized to control (+R) are shown. N = 3 independent experiments. Mean ± SEM is shown; two-tailed unpaired Student’s *t*-test; ns: not significant; *: *p*<0.05; **: *p*<0.01; ***: *p*<0.005; R: arginine; *le*: long exposure; -R +R: arginine depletion followed by re-supply.

During DNA replication, PCNA is repeatedly loaded and unloaded by the replicating clamp loader, replication factor C (RFC) complex, and an alternative PCNA ring opener, ATAD5 (ELG1 in yeast)-RFC-like complex (ATAD5-RLC). ATAD5-RLC unloads replication-coupled PCNA (Arbel et al., 2021). Maintaining the balance of PCNA loaded onto nascent DNA strands is crucial for DNA replication (Choe and Moldovan, 2017). To further test the effect of arginine shortage on chromatin-bound PCNA levels, we performed chromatin fractionation to determine the chromatin-associated fraction of subunits of the PCNA unloader, RLC (ATAD5/RFC2-5) complex (Arbel et al., 2021). Consistent with the increased PCNA levels, the chromatin-associated ATAD5, RFC2, and RFC5, subunits of the RLC complex, were significantly reduced in arginine-deprived MDA-MB-231 cells. In addition, the total level of these proteins as shown in whole-cell extracts was also reduced upon arginine removal (**Fig. S6B**). A similar observation was made in arginine-deprived BT-549 cells except that ATAD5 level was almost undetectable (**Fig. S6B**). While consistent with the reduced H4K5Ac and H4K12Ac deposition, a marked reduction in chromatin-bound BRD4, an acetyl-histone-binding chromatin reader (revised **Fig. S6B**) seems to conflict with increased PCNA loading. A previous study showed that ATAD5, the major PCNA unloader, had dominant effects on PCNA unloading, and overexpression of BRD4 significantly induced PCNA accumulation on chromatin. However, depletion of BRD4 can only slightly decrease the levels of PCNA on nascent DNA (Kang et al., 2019). Therefore, even though the downregulation of BRD4 should promote PCNA unloading during arginine shortage, the decreased levels of ATAD5, RFC2, and RFC5 could eventually lead to insufficient unloading of PCNA. Together, these findings suggest that the reduced chromatin binding of the PCNA-unloader ATAD5-RLC causes the increased retention of PCNA in the arginine-deprived cells. It remains to be elucidated how arginine removal affects the crosstalk among BRD4, ATAD5 and H4K5ac/H4K12ac to fine-tune PCNA retention on nascent DNA strands.

Given that altered ubiquitination of replication machinery is a hallmark of stressed replication forks (Nakamura et al., 2021), we investigated if elevated levels of PCNA might correspond to altered PCNA ubiquitination. PCNA can be either monoubiquitinated or polyubiquitinated to activate translesion synthesis or fork reversal (Hoege et al., 2002; Kannouche et al., 2004). To test whether PCNA, is ubiquitinated upon arginine removal in a manner similar to that observed under canonical replication stress, HEK-293T cells stably expressing FLAG-tagged PCNA were transiently transfected with Myc-tagged ubiquitin in our biochemical analyses. Following arginine removal (4 h) and immunoprecipitation with an anti-FLAG antibody, an increased K63-linked polyubiquitination and K164 monoubiquitination of PCNA was detected (**Fig. 5C**). HLTF is one of the E3 ligases which catalyzes PCNA K63-linked polyubiquitination in a RAD6-RAD18-dependent manner (Bai et al., 2020; Kile et al., 2015). Immunoprecipitation followed by Western analyses further revealed that arginine removal enhanced the recruitment of RAD6, RAD18 and HLTF to PCNA (**Fig. 5D**). The replenishment of arginine reversed arginine shortage-increased K63-linked polyubiquitination and recruitment of RAD6, RAD18, and HLTF to PCNA without affecting PCNA levels (**Fig. 5E**). Lastly, knockdown of HLTF abolished arginine shortage-induced K63-linked polyubiquitination of PCNA (**Fig. 5F**). Next, we performed a chromatin fractionation and found the levels of chromatin-bound PCNA, HLTF, RAD18, and RAD6 were all significantly higher in arginine-deprived cells, indirectly supporting that increased associations among RAD6, RAD18, HLTF and PCNA on chromatin resulted in more PCNA K63-linked polyubiquitination at forks (**Fig. S6A**). Based on these observations, we propose that HLTF-mediated K63-linked polyubiquitination of PCNA and/or its associated proteins increases in response to extracellular arginine shortage in a reversible manner. To examine the possibility whether other K63-linked E3 ligases, such as SHPRH (Bruhl et al., 2019), is involved in arginine shortage-induced PCNA K63-linked polyubiquitination, SHPRH was knocked down in arginine-deprived cells (**Fig. S6C**). However, depletion of SHPRH failed to reverse arginine shortage-induced K63-linked polyubiquitination. These findings demonstrate that shortage of arginine induces HLTF-, but not SHPRH-, mediated K63-linked polyubiquitination of PCNA. To examine whether there is a link between histone H4 synthesis and PCNA K63-linked polyubiquitination, we impaired histone H4 biosynthesis by knocking down FLASH or SLBP, both are essential for histone biosynthesis (Mejlvang et al., 2014) (**Fig. S6D**). Indeed, knockdown of FLASH or SLBP decreased newly synthesized H4 marks, as previously reported, and increased K63-linked polyubiquitination of PCNA (**Fig. S6D**). This observation links arginine shortage-suppressed *de novo* histone H4 synthesis to HLTF-mediated PCNA K63-ubiquitination.

### Arginine shortage promotes HLTF-dependent genotoxin adaption

To mechanistically explore the role of HLTF-mediated K63-linked polyubiquitination of PCNA and/or its associated proteins in response to arginine shortage, we first sought to analyze how HLTF-loss altered cellular responses to arginine removal. Previously, HLTF-loss reportedly led to an unrestrained fork progression (Bai et al., 2020; Kile et al., 2015). To confirm and extend this finding in arginine-starved cells, we used two different siRNAs to knock down HLTF and monitored fork progression under ± arginine in MDA-MB-231 cells as described in **Fig. 1C**. Compared to siCtrl-treated cells, knockdown of HLTF alone did not markedly affect the level of H4K5ac and H4K12ac marks (**Fig. S7A**) and the ratio of IdU/CldU track (**Fig. 6A**) in unstarved cells, respectively. However, knockdown of HLTF partially rescued the arginine shortage-suppressed IdU/CldU ratio (**Fig. 6A**). To provide additional evidence supporting the effect of PCNA ubiquitination on fork slowing in arginine-starved cells, isogenic paired HEK-293T-WT and HEK-293T-K164R cells (Thakar et al., 2020) were employed. **Fig. 6B** shows that the PCNA K164R mutation, at least in part, rescued IdU/CldU ratio in arginine-starved cells and inactivating HLTF in the K164R cells had no further effect. Consistent with the idea that HLTF has an activity of slowing fork progression in cells experiencing replication stress (Kile et al., 2015), our observation confirmed that HLTF restrains replication fork progression in arginine-starved cells.

**Fig. 6.**
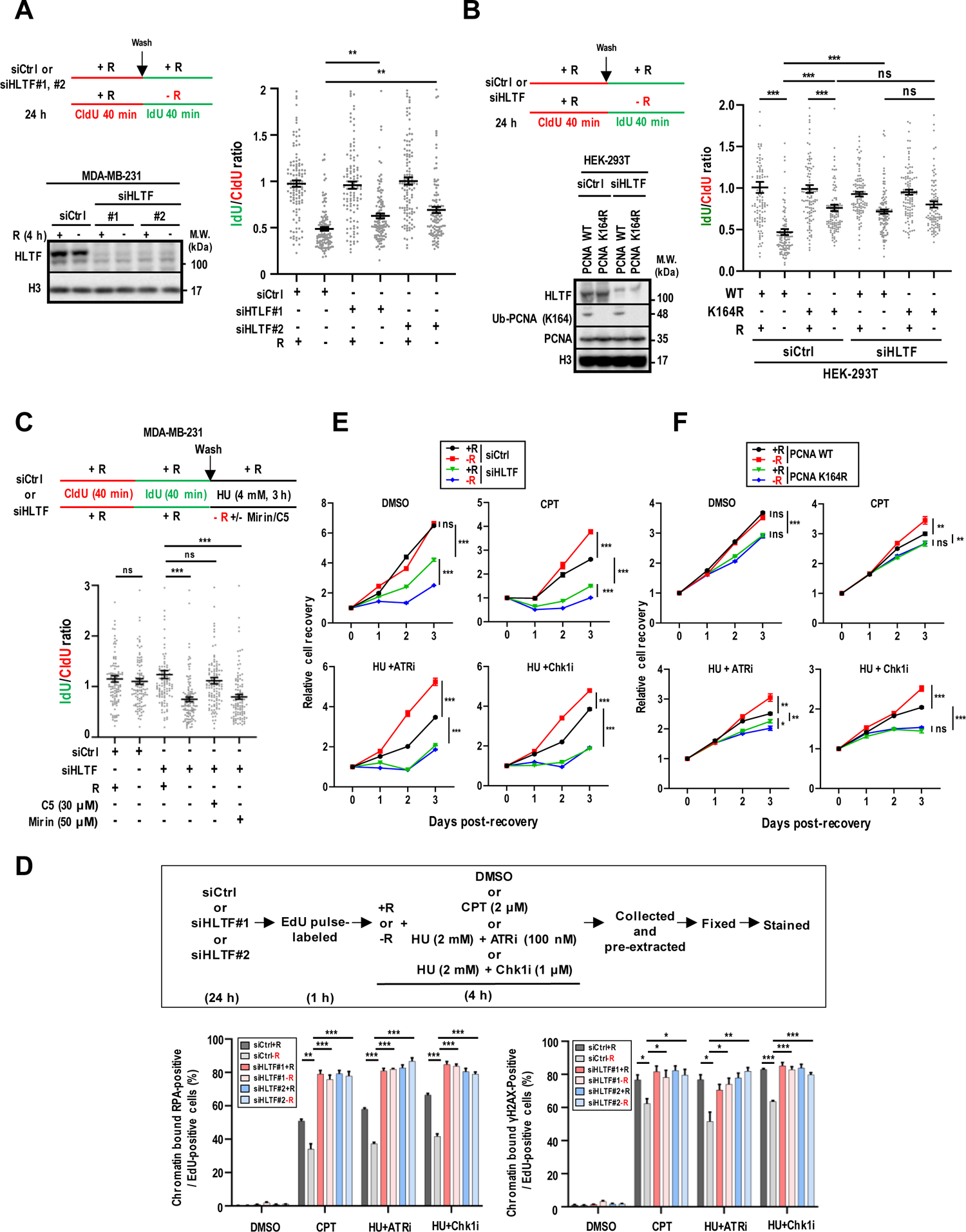
HLTF and PCNA protects genome stability in arginine-starved cells. (**A**) Analysis following MDA-MB-231 cells treated with indicated siRNAs targeting HLTF and ± arginine. *Top left*: Experimental outline. *Bottom left:* Representative Western blot analysis of HLTF. *Right*: Quantitation of IdU/CldU ratio. N = 3 independent experiments. Each dot represents one fiber. At least 100 tracts were analyzed per sample. (**B**) Analysis of HEK-239T cells overexpressing wild-type (WT) or mutated (K164R) PCNA treated with siRNA targeting HLTF ± arginine. *Top left*: Experimental outline. *Bottom left:* Representative Western blot analysis of K164 ubiquitinated PCNA (ub-PCNA) and HLTF. *Right*: Quantitation of IdU/CldU ratio. N = 3 independent experiments. Each dot represents one fiber. At least 100 tracts were analyzed per sample. (**C**) Analysis of the effect of DNA2 inhibitor C5 or MRE11 inhibitor mirin on replication fork stability under ± arginine and exposure to HU in HLTF-knockdown MDA-MB-231 cells. *Top:* Experimental outline. *Bottom:* Quantification of IdU/CldU ratio. N = 3 independent experiments. Each dot represents one fiber. At least 100 tracts were analyzed per sample. (**D**) Analysis of HLTF-knockdown on chromatin-bound RPA and γH2AX in MDA-MB-231 cells grown under ± arginine and subjected to genotoxic insult. *Top*: Experimental outline. *Bottom*: Quantification of chromatin-bound RPA^+^ or γH2AX^+^ cells following flow analysis. At least 10,000 events were collected and analyzed. N = 3 independent exp€ments. (**E**) Relative cell recovery of control or HLTF-knockdown MDA-MB-231 cells after treatment with indicated genotoxic agents and ± arginine. N= 3 independent experiments. (**F**) Relative cell recovery of wild-type (WT) or mutated (K164R) PCNA-expressing HEK-293T cells after treatment with indicated genotoxic agents and ± arginine. N = 3 independent experiments. Mean ± SEM is shown; ns: not significant; *: *p*<0.05; **: *p*<0.01; ***: *p*<0.005; two-tailed unpaired Student’s *t*-test; R: arginine.

Next, we sought to address the contribution of HLTF to the integrity of arginine shortage-stressed replication forks. In order to do this, we treated HLTF-depleted MDA-MB-231 cells with a high concentration of HU (4 mM), which has been shown to promote fork reversal (Zellweger et al., 2015), and measured the IdU/CldU-labeled fiber ratio under ± arginine conditions (**Fig. 6C**). siRNA depletion of HLTF alone had no effect on fork degradation in cells cultured with HU (4 mM) and arginine. In contrast, the removal of arginine and exposure to HU (4 mM) notably accelerated fork degradation in HLTF-depleted cells (**Fig. 6C**, *lane #4 vs. lane #3*). Collectively, these findings suggest that HLTF is critical to protect nascent DNA from degradation in arginine-deprived cells when exposed to HU (4 mM). Mirin, an MRE11 inhibitor, did not suppress nascent strand degradation in arginine-starved, HLTF-depleted cells (**Fig. 6C**). In contrast, treatment with C5, an inhibitor of the nuclease DNA2, restored the nascent track length (**Fig. 6C**), indicating the possible involvement of DNA2-mediated nascent DNA nucleolysis in arginine-starved, HLTF-depleted cells. To determine whether the protective function of HLTF relies on its Hiran domain to bind specifically to 3’-end of ssDNA as previously reported, we tested the fork integrity in cells overexpressing siRNA-resistant wild-type HLTF and Hiran-mutated (R71E) HLTF (Taglialatela et al., 2017). **Fig. S7B** shows that knockdown of endogenous HLTF in MDA-MB-231 cells induced the degradation of replication fork; however, both wild-type and Hiran-mutated HLTF can almost fully restore nascent DNA strands (**Fig. S7B**), implicating that HLTF exerts its protective role in arginine-depleted cells independent of its ssDNA binding ability. To further investigate whether other known translocases share the protective function of HLTF, the fork integrity was tested in SMARCAL1 (Kolinjivadi et al., 2017)- or ZRANB3 (Vujanovic et al., 2017)-knockdown cells (**Fig. S7C**). Results showed that knockdown of SMARCAL1 or ZRANB3 did not induce degradation of replication fork in arginine-starved cells (**Fig. S7C**). These observations indicated that HLTF-dependent fork reversal is not involved in resolving arginine shortage-induced fork stalling.

To determine the impact of HLTF on DNA-damage signaling upon arginine shortage, we then used a flow cytometry-based method (Forment and Jackson, 2015) to quantitatively monitor RPA and phospho-H2AX or γH2AX chromatin binding in S-phase cells (**Fig. 6D**, *upper diagram*). As expected, brief exposure to high concentration of CPT (2 μM, 4 h) increased both RPA and γH2AX chromatin binding in MDA-MB-231 S-phase cells cultured in full medium and arginine shortage attenuated both signals (**Fig. 6D**, *lower panels*). Notably, HLTF-loss increased both RPA and γH2AX chromatin binding in EdU-positive cells under arginine shortage (**Fig. 6D**, *lower panels, lanes 10, 12 vs. lane 8*). A similar trend was noted (**Fig. 6D**, *lower panels*) in cells treated with HU+ATRi *(lanes 16, 18 vs. lane 14)* or HU+CHK1i *(lanes 22, 24 vs. lane 20)* under arginine shortage. Lastly, we determined the role of HLTF in protecting MDA-MB-231 cells from genotoxins under ± arginine conditions. Compared to **Fig. 4E**, HLTF-knockdown mitigated the advantage conferred by arginine removal on cell recovery from indicated genotoxic treatments (**Fig. 6E**). Similar observations, albeit less pronounced, were made in PCNA-mutated (K164R) HEK-293T cells (**Fig. 6F**). While PCNA K63-linked polyubiquitination is likely associated with poor recovery of HLTF-knockdown cells from CPT, HU+ATRi, or HU+Chk1i, the exact contribution to the better recovery remains unclear. It is possible that HLTF-catalyzed PCNA K164 ubiquitination (**Fig. 5F**) is indispensable for arginine shortage to enable cells to better cope with DNA damage induced by a variety of genotoxic agents.

AOH1996 is the lead compound targeting the L126-Y133 region of PCNA to interfere with its interaction with partners (*Gu et al., 2019*). To address PCNA function in further detail, we determined the effect of AOH1996 on the fork integrity in arginine-deprived cells. aniPOND assays show that AOH1996 suppressed PCNA accumulation on replication forks (**Fig. S8A**) without affecting the levels of newly synthesized H4 marks (**Fig. S8B**). Like HLTF-knockdown, a fork protection assay showed that AOH1996 accelerated fork degradation in arginine-deprived cells (**Fig. S8C**, *lane 4 vs. lane 3*). Next, to address whether AOH1996 could reverse arginine shortage-induced K63-linked polyubiquitination of PCNA and genotoxin resistance, immunoprecipitation, a flow cytometry-based assay (Forment and Jackson, 2015) and a cell recovery assay were performed. We showed that AOH1996 reversed arginine shortage-induced K63-linked polyubiquitination of PCNA (**Fig. S8D**, *lane 5 vs. lane 3*). AOH1996 also increased γH2AX accumulation on chromatin in arginine-starved, EdU-positive cells treated with genotoxins (**Fig. S8E**). Additionally, AOH1996 abolished the protection provided by arginine shortage on cells recovered from genotoxic insult (**Fig. S8F**). Taken together, we suggest that PCNA loading, and its ubiquitination are important for maintaining fork integrity and recovery from genotoxicity in arginine-deprived cells. Collectively, we propose a model, based on our findings, that arginine shortage inhibits H4 translation to stall DNA replication and enables HLTF-mediated K63-linked polyubiquitination of PCNA and/or its associated proteins and hyper-PCNA accumulation on nascent DNA to protect arginine-depleted cells against genotoxins (**Fig. S9**).

## DISCUSSION

We identified arginine, a nutrient that is frequently limited within the TME (Pan et al., 2016), as a DNA replication modulator. We find that arginine shortage significantly impacts the progression of replication forks by inhibiting histone H4 translation and promoting the hyper-accumulation and HLTF-mediated ubiquitination of PCNA on nascent DNA strands. Significantly, our results link arginine shortage to genotoxic adaptation.

Replication-dependent histone biosynthesis has been generally regarded as a critical regulator for DNA replication and cell cycle progression (Armstrong and Spencer, 2021; Gunesdogan et al., 2014; Mejlvang et al., 2014). We find that increased abundance of histone H4 alone is sufficient to promote DNA replication elongation under extracellular arginine shortage. Arginine accounts for, on average, approximately 4.2% of the amino acid residues within known vertebrate protein sequences, which makes it neither a particularly common nor a particularly rare amino acid. Although all four of the core histone amino acid sequences contain a higher-than-average number of arginine residues, only histone H4 protein translation is markedly sensitive to acute arginine shortage, leading to a shutoff of DNA replication. Darnell et al. reported the selective loss of arginine tRNA charging during arginine shortage regulates translation via ribosome pausing at specific arginine codons (Darnell et al., 2018). In line with this, there is a higher percentage of the putative arginine pause-site codons (CGC and CGU) in histone H4 (10.6%) than those in histone H3 (8.8%) and histone H2B (6.3%), respectively (**Table 1-3**). Altogether, our observations raise the possibility that arginine residues in histone H4 may carry out a nutrient-sensing function by changing its synthesis in response to extracellular arginine availability. Finally, our work suggests that extracellular arginine shortage deaccelerates DNA replication independent of arginine *de novo* synthesized from urea cycle and ASS1 abundance. This phenomenon warrants further investigation to identify and characterize the usages of different sources of arginine in multiple biological pathways. For example, whether and how limitation in extracellular arginine supply affects pools tRNA^Arg^ CGC and CGU isoacceptors more than endogenous arginine.

**Table 1.** The mRNA and amino acids composition of *Homo sapiens* histone H4 (H4C1 H4).

|  |  |  |  |  |  |  |  |  |  |  |  |  |  |  |  |  |  |  |  |  |
| --- | --- | --- | --- | --- | --- | --- | --- | --- | --- | --- | --- | --- | --- | --- | --- | --- | --- | --- | --- | --- |
| **mRNA** | AUG | UCU | GGA | CGU | GGU | AAG | GGC | GGG | AAG | GGU | UUG | GGU | AAG | GGG | GGU | GCC | AAG | CGC | CAC | CGC |
| **Protein** | MET | SER | GLY | ARG | GLY | LYS | GLY | GLY | LYS | GLY | LEU | GLY | LYS | GLY | GLY | ALA | LYS | ARG | HIS | ARG |
|  | AAG | GUG | UUG | CGU | GAC | AAC | AUC | CAG | GGC | AUC | ACC | AAG | CCG | GCC | AUC | CGG | CGU | CUG | GCC | CGG |
|  | LYS | VAL | LEU | ARG | ASP | ASN | ILE | GLN | GLY | ILE | THR | LYS | PRO | ALA | ILE | ARG | ARG | LEU | ALA | ARG |
|  | CGU | GGC | GGU | GUG | AAG | CGG | AUC | UCU | GGU | CUG | AUC | UAC | GAG | GAG | ACU | CGC | GGG | GUG | CUC | AAG |
|  | ARG | GLY | GLY | VAL | LYS | ARG | ILE | SER | GLY | LEU | ILE | TYR | GLU | GLU | THR | ARG | GLY | VAL | LEU | LYS |
|  | GUG | UUU | UUG | GAG | AAC | GUG | AUC | CGU | GAC | GCU | GUC | ACC | UAU | ACG | GAG | CAC | GCC | AAG | CGC | AAG |
|  | VAL | PHE | LEU | GLU | ASN | VAL | ILE | ARG | ASP | ALA | VAL | THR | TYR | THR | GLU | HIS | ALA | LYS | ARG | LYS |
|  | ACA | GUC | ACU | GCC | AUG | GAC | GUG | GUC | UAC | GCG | CUU | AAG | CGC | CAG | GGA | CGC | ACC | CUU | UAU | GGC |
|  | THR | VAL | THR | ALA | MET | ASP | VAL | VAL | TYR | ALA | LEU | LYS | ARG | GLN | GLY | ARG | THR | LEU | TYR | GLY |
|  | UUU | GGC | GGU | UAA |  |  |  |  |  |  |  |  |  |  |  |  |  |  |  |  |
|  | PHE | GLY | GLY | STOP |  |  |  |  |  |  |  |  |  |  |  |  |  |  |  |  |

(Ribosome pause-site arginine codons: CGU, CGC; non-ribosome pause-site arginine codons: CGG, CGA, AGA, AGG)(Ribosome pause-site encoded arginine: ARG; non-ribosome pause-site encoded arginine: ARG)
Arginine pausing codon/all codons: 11/104 (10.58%)
CGU/all codons: 5/104 (CGU/arginine codon: 5/14; CUG/ribosome pausing arginine codon: 5/11)
CGC/all codons: 6/104 (CGC/arginine codon 6/14; CGG/ribosome pausing arginine codon: 6/11)

**Table 2.** The mRNA and amino acids composition of *Homo sapiens* histone H3 (H3C14).

|  |  |  |  |  |  |  |  |  |  |  |  |  |  |  |  |  |  |  |  |  |
| --- | --- | --- | --- | --- | --- | --- | --- | --- | --- | --- | --- | --- | --- | --- | --- | --- | --- | --- | --- | --- |
| **mRNA** | AUG | GCC | CGU | ACU | AAG | CAG | ACU | GCU | CGC | AAG | UCG | ACC | GGC | GGC | AAG | GCC | CCG | AGG | AAG | CAG |
| **Protein** | MET | ALA | ARG | THR | LYS | GLN | THR | ALA | ARG | LYS | SER | THR | GLY | GLY | LYS | ALA | PRO | ARG | LYS | GLN |
|  | CUG | GCC | ACC | AAG | GCG | GCC | CGC | AAG | AGC | GCG | CCG | GCC | ACG | GGC | GGG | GUG | AAG | AAG | CCG | CAC |
|  | LEU | ALA | THR | LYS | ALA | ALA | ARG | LYS | SER | ALA | PRO | ALA | THR | GLY | GLY | VAL | LYS | LYS | PRO | HIS |
|  | CGC | UAC | CGG | CCC | GGC | ACC | GUA | GCC | CUG | CGG | GAG | AUC | CGG | CGC | UAC | CAG | AAG | UCC | ACG | GAG |
|  | ARG | TYR | ARG | PRO | GLY | THR | VAL | ALA | LEU | ARG | GLU | ILE | ARG | ARG | TYR | GLN | LYS | SER | THR | GLU |
|  | CUG | CUG | AUC | CGC | AAG | CUG | CCC | UUC | CAG | CGG | CUG | GUA | CGC | GAG | AUC | GCG | CAG | GAC | UUU | AAG |
|  | LEU | LEU | ILE | ARG | LYS | LEU | PRO | PHE | GLN | ARG | LEU | VAL | ARG | GLU | ILE | ALA | GLN | ASP | PHE | LYS |
|  | ACG | GAC | CUG | CGC | UUC | CAG | AGC | UCG | GCC | GUG | AUG | GCG | CUG | CAG | GAG | GCC | AGC | GAG | GCC | UAC |
|  | THR | ASP | LEU | ARG | PHE | GLN | SER | SER | ALA | VAL | MET | ALA | LEU | GLN | GLU | ALA | SER | GLU | ALA | TYR |
|  | CUG | GUG | GGG | CUG | UUC | GAA | GAC | ACG | AAC | CUG | UGC | GCC | AUC | CAC | GCC | AAG | CGC | GUG | ACC | AUU |
|  | LEU | VAL | GLY | LEU | PHE | GLU | ASP | THR | ASN | LEU | CYS | ALA | ILE | HIS | ALA | LYS | ARG | VAL | THR | ILE |
|  | AUG | CCC | AAG | GAC | AUC | CAG | CUG | GCC | CGC | CGC | AUC | CGU | GGA | GAG | CGG | GCU | UAA |  |  |  |
|  | MET | PRO | LYS | ASP | ILE | GLN | LEU | ALA | ARG | ARG | ILE | ARG | GLY | GLU | ARG | ALA | STOP |  |  |  |

(Ribosome pause-site arginine codons: CGU, CGC; non-ribosome pause-site arginine codons: CGG, CGA, AGA, AGG)(Ribosome pause-site encoded arginine: ARG; non-ribosome pause-site encoded arginine: ARG)
Arginine pausing codon/all codons: 12/137 (8.76%)
CGU/all codons: 2/137 (CGU/arginine codon: 2/18; CUG/ribosome pausing arginine codon: 2/12)
CGC/all codons: 10/137 (CGC/arginine codon 10/18; CGG/ribosome pausing arginine codon: 10/12)

**Table 3.**
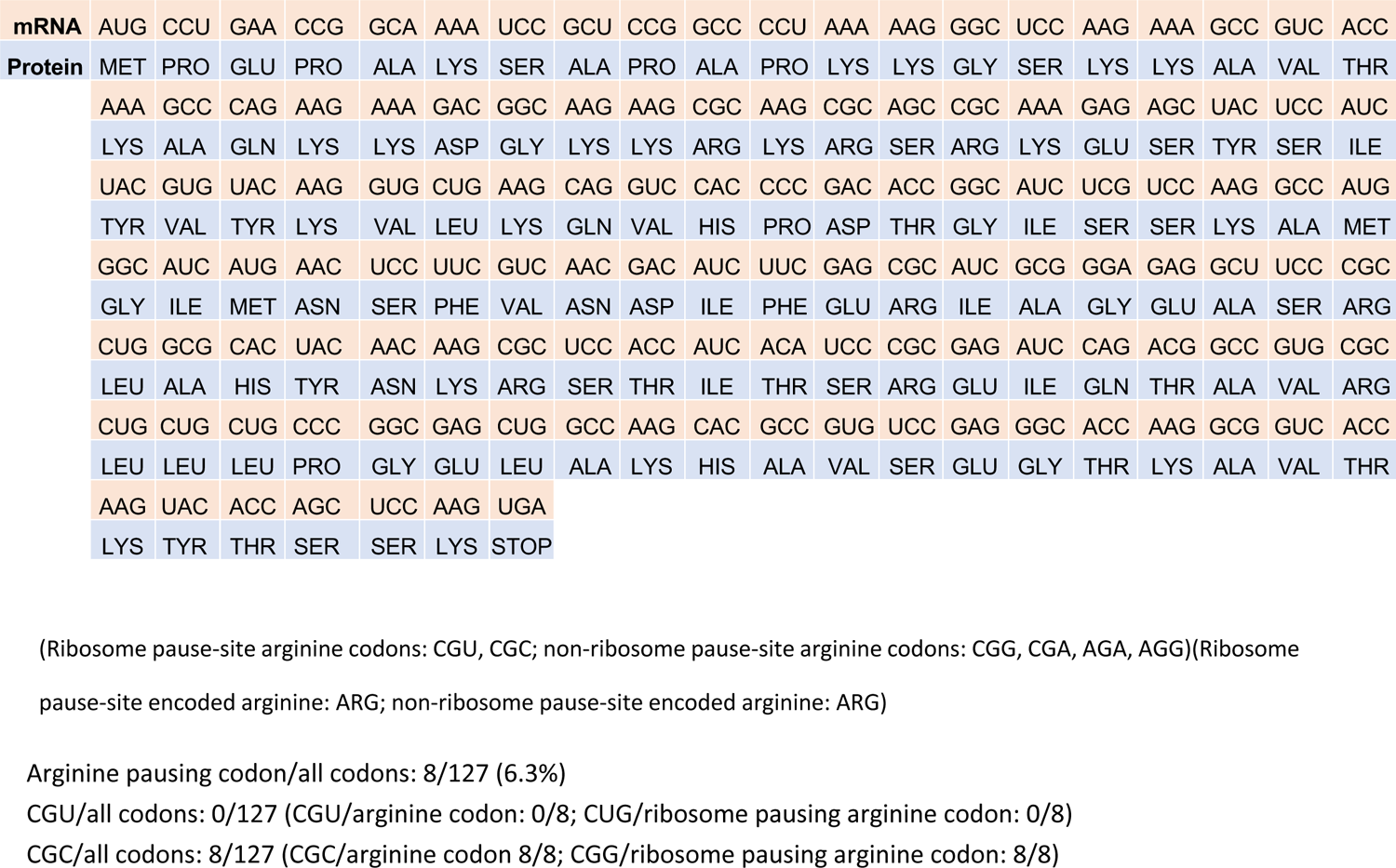
The mRNA and amino acids composition of *Homo sapiens* histone H2 (H2BC21).

| mRNA | AUG | CCU | GAA | CCG | GCA | AAA | UCC | GCU | CCG | GCC | CCU | AAA | AAG | GGC | UCC | AAG | AAA | GCC | GUC | ACC |
| --- | --- | --- | --- | --- | --- | --- | --- | --- | --- | --- | --- | --- | --- | --- | --- | --- | --- | --- | --- | --- |
| Protein | MET | PRO | GLU | PRO | ALA | LYS | SER | ALA | PRO | ALA | PRO | LYS | LYS | GLY | SER | LYS | LYS | ALA | VAL | THR |
|  | AAA | GCC | CAG | AAG | AAA | GAC | GGC | AAG | AAG | CGC | AAG | CGC | AGC | CGC | AAA | GAG | AGC | UAC | UCC | AUC |
|  | LYS | ALA | GLN | LYS | LYS | ASP | GLY | LYS | LYS | ARG | LYS | ARG | SER | ARG | LYS | GLU | SER | TYR | SER | ILE |
|  | UAC | GUG | UAC | AAG | GUG | CUG | AAG | CAG | GUC | CAC | CCC | GAC | ACC | GGC | AUC | UCG | UCC | AAG | GCC | AUG |
|  | TYR | VAL | TYR | LYS | VAL | LEU | LYS | GLN | VAL | HIS | PRO | ASP | THR | GLY | ILE | SER | SER | LYS | ALA | MET |
|  | GGC | AUC | AUG | AAC | UCC | UUC | GUC | AAC | GAC | AUC | UUC | GAG | CGC | AUC | GCG | GGA | GAG | GCU | UCC | CGC |
|  | GLY | ILE | MET | ASN | SER | PHE | VAL | ASN | ASP | ILE | PHE | GLU | ARG | ILE | ALA | GLY | GLU | ALA | SER | ARG |
|  | CUG | GCG | CAC | UAC | AAC | AAG | CGC | UCC | ACC | AUC | ACA | UCC | CGC | GAG | AUC | CAG | ACG | GCC | GUG | CGC |
|  | LEU | ALA | HIS | TYR | ASN | LYS | ARG | SER | THR | ILE | THR | SER | ARG | GLU | ILE | GLN | THR | ALA | VAL | ARG |
|  | CUG | CUG | CUG | CCC | GGC | GAG | CUG | GCC | AAG | CAC | GCC | GUG | UCC | GAG | GGC | ACC | AAG | GCG | GUC | ACC |
|  | LEU | LEU | LEU | PRO | GLY | GLU | LEU | ALA | LYS | HIS | ALA | VAL | SER | GLU | GLY | THR | LYS | ALA | VAL | THR |
|  | AAG | UAC | ACC | AGC | UCC | AAG | UGA |  |  |  |  |  |  |  |  |  |  |  |  |  |
|  | LYS | TYR | THR | SER | SER | LYS | STOP |  |  |  |  |  |  |  |  |  |  |  |  |  |

(Ribosome pause-site arginine codons: CGU, CGC; non-ribosome pause-site arginine codons: CGG, CGA, AGA, AGG)(Ribosome pause-site encoded arginine: ARG; non-ribosome pause-site encoded arginine: ARG)
Arginine pausing codon/all codons: 8/127 (6.3%)
CGU/all codons: 0/127 (CGU/arginine codon: 0/8; CUG/ribosome pausing arginine codon: 0/8)
CGC/all codons: 8/127 (CGC/arginine codon 8/8; CGG/ribosome pausing arginine codon: 8/8)

Besides the extracellular arginine-histone H4 link, our studies also strongly suggest that an arginine-PCNA link is essential for sustaining DNA replication as well (**Fig. S9**). At this point, we cannot exclude that additional factor, such as Chromatin Assembly Factor-1 (CAF-1 (Smith and Stillman, 1989)) or other histone chaperones that are recruited to the replisome (*i.e*., FACT and ASF1 (Alabert et al., 2017)), may function on connecting arginine-dependent H4 translation with HLTF-catalyzed PCNA K63-linked polyubiquitination on nascent strands to impair chromatin assembly, nor can we exclude the possibility that arginine availability may affect the interaction between PCNA and other key components involved in DNA replication (*i.e*., DNA polymerases, replication factor C (RFC) and RFC-like complexes). All these proteins coordinate the recycling of histones or PCNA to spatially maintain the landscape of histone post-translational modifications or PCNA occupancy. Indeed, newly replicated DNA is readily assembled into chromatin, a process that constitutes the first critical step to the re-establishment of epigenetic modifications on histones genome wide (MacAlpine and Almouzni, 2013) In contrast, we do not observe evidence for the occurrence of large ssDNA tracts during arginine shortage but cannot exclude DNA nicks as PCNA interacts with DNA ligase I (Choe and Moldovan, 2017). Taken together, arginine shortage results in increased PCNA K63-linked polyubiquitination, which has the potential to serve as a marker for DNA replication stress. Needless to say, additional studies are needed to provide definitive evidence for the role of arginine in regulating PCN’’s residence on the nascent DNA strands or genome stability maintenance.

An open question pertains to the notion that PCNA was the only protein that significantly accumulated on the nascent DNA strands in arginine-starved cells. The technical aspects such as the efficiency of cross-linking or the stringency of the conditions used for chromatin purification might have caused the loss of loosely bound PCNA-associated factors. One surprising conclusion from these studies is that arginine shortage activates Rad6/Rad18-mediated PCNA K164 monoubiquitination and HLTF-catalyzed PCNA K63-linked polyubiquitination. Considering that PCNA needs to be recycled repeatedly during replication in order for DNA replication to progress continuously (Shibahara and Stillman, 1999), it is interesting to find that arginine plays a role in regulating the timely PCNA unloading from nascent DNA strands by controlling the levels of proteins comprising the unloading machinery. Whether HLTF-mediated K63-linked polyubiquitination of PCNA and/or associated proteins prevents the dissociation of PCNA from the nascent DNA strands is sufficient to slow DNA replication remains to be determined. We surmise that the K63-linked E3 ligase activity of HLTF, but not SHPRH, is crucial to prolong the association of PCNA on nascent strands and potentially alter interactions between PCNA and its partners. Alternatively, K63-linked, polyubiquitinated PCNA at newly synthesized DNA stand can block the sliding of PCNA molecules on DNA, resulting in the accumulation of PCNA behind the stalled replication fork in arginine-starved cells. In any case, it will be interesting to further investigate the regulation of HLTF activity and its possible interplay with DNA replication machinery: what is the role of K63-linked polyubiquitination of PCNA and/or associated proteins in deaccelerating fork progression or enabling cells to survive genotoxins? How is PCNA unloading coordinated with histone deposition in the wake of arginine shortage during DNA replication? Given the important function of PCNA in DNA replication and DNA damage repair, how is its activity regulated by arginine to safeguard genome stability?

Finally, we still do not fully understand the impact of arginine shortage on genome integrity maintenance. The S-phase arrest observed in arginine-starved cells alone appears insufficient to explain the reduced genotoxic damage sensitivity, in particular, arginine shortage only slows DNA replication without activating a damage signal. In our report, the consequences of HLTF-mediated genome protection seems to occur in an arginine-dependent manner, possibly through the recruitment of interacting partners or its E3 ligase activity. Peng et al. have shown that long-term HU (4 mM) treatment can result in fork degradation in FANCJ-knockout cells, while HLTF depletion significantly reduces fork degradation in FANCJ-knockout cells (Peng et al., 2018). However, we find that the loss-of-HLTF accelerates HU (4 mM)-induced fork degradation in arginine-starved cells. In addition, K63-linked, polyubiquitinated PCNA blocks DNA2 from loading to replication forks (Thakar et al., 2020). This is consistent with our observations that arginine shortage results in DNA2-dependent but MRE11-independent nucleolytic degradation of nascent DNA at stalled replication forks in HLTF-depleted cells (**Fig. 6C**). Moreover, HLTF has been shown to lead to resistance to DNA damaging agents in multiple studies, which is in consistent with our data here that the loss of HLTF is shown to sensitize arginine-deprived cells (Blastyak et al., 2010; Cho et al., 2011; Unk et al., 2008; Ye et al., 2015). In general, HLTF has been predominantly known for its functions in post-replication repair, where it stimulates replication fork reversal to bypass the blocks and restart replication forks in response to replication stress (Quinet et al., 2021). Recently, van Toorn et al. reported that HLTF is required for efficient nucleotide excision repair (van Toorn et al., 2022). In our study, we ruled out the involvement of fork reversal by HLTF and other translocases in managing the replication stress induced by arginine shortage (**Fig. S7B, S7C**). However, the detailed mechanism of how HLTF-mediated K63-linked polyubiquitinated PCNA maintains genome integrity independent of fork reversal under arginine shortage remains unclear and is worth further investigation.

In summary, we have uncovered a hitherto unidentified pathway, the histone H4 translation-arrest and HLTF-mediated PCNA K63-linked polyubiquitination, which provides an instant and reversible response, slowing DNA replication, when extracellular arginine availability is altered (**Fig. S9**). Our model predicts that endogenous arginine synthesized by ASS1 is not sufficient to drive DNA replication. Mechanisms that modulate DNA replication, elongation and prevent replication fork disruption might be more widespread than we recognized. Our results also suggest that the instant replication fork slowing in response to extracellular arginine shortage helps maintain genome stability against a variety of agents by preventing deleterious consequences from occurring in proliferating cells. The arrested replication will provide an opportunity for lesions, if acquired, to be repaired, and replication to resume with fewer obstacles after arginine becomes available. The H4 translation inhibition- and PCNA K63-linked polyubiquitination-mediated arrest that prevents genotoxin-induced genomic instability during transient regional arginine shortage, in turn, corroborates that local nutrient availability within the TME which promotes heterogeneity between tumor cells residing in proximity or not to the vasculature. Our study also makes a clear point that arginine shortage provides a protective effect against genotoxicity. Presumably, this protective effect would be a common phenomenon naturally occurring in the core regions of various types of solid tumors regardless of its genetic, epigenetic, and metabolic backgrounds. A recent report showed that administration of arginine sensitizes tumors to radiation therapy (Marullo et al., 2021), which supports our assumption that arginine levels should be taken into consideration when developing therapeutic approaches and might significantly impact the response to anti-cancer therapy.

### Limitations of the study

Our study identifies extracellular arginine as a key factor that stimulates DNA replication fork progression through promotion of newly synthesized histone H4 for chromatin assembly on nascent DNA strands and facilitation of PCNA recycling. Based on our protein histone H4 transfection data and the involvement of the HLTF-mediated PCNA K63-linked polyubiquitination, we favor a model in which arginine shortage acts at the level of newly synthesized histone H4 supply to subsequently cause PCNA accumulation on nascent strands. One caveat is that the lack of newly synthesized histone H3 could theoretically slow down the DNA replication fork and accumulate PCNA on nascent strands. Keeping the limitation of the immunoprecipitation assay in mind, we were not able to determine whether arginine shortage promotes the K63-linked polyubiquitination of chromatin-bound PCNA in an unequivocal manner. Determining the role of ZRANB3 in the context of arginine shortage in more detail remains of importance to fully understand the mechanistic details of the transition of unmodified PCNA to K63-linked polyubiquitination state, which protects the integrity of replication forks. Lastly, our studies also pave the way for future investigations in understanding how arginine-dependent newly synthesized histone H4 deposition pathways are integrated during DNA replication.

## Acknowledgements

We thank Drs. Mark LaBarge, Lei Jiang, Shao-Chun Wang, Anita K. Hopper, Chathurani S. Jayasena and members of Dr. Ann’s, Stark’s and Kung’s laboratories for helpful discussion on the manuscript. Dr. Glenn Manthey for editing the manuscript. Immunohistochemistry staining was performed by Pathology Core of City of Hope by Dr. Aimin Li. This work is supported by funds from the National Institutes of Health (R01CA220693 to DKA, R01CA197506 to JMS, R01GM134681 to GLM, T32CA186895 to AAK, T32CA221709-01 for AK and P30CA33572 for City of Hope Core Facilities). The content is solely the responsibility of the authors and does not necessarily represent the official views of the National Institutes of Health.

## Author contributions

YCW, JMS, HJK and DKA devised the study. YCW performed most of the experiments and analysis. AAK contributed to DNA combing assay. AK contributed to the development of newly synthesized protein assays. YHC and WKC contributed to subcellular fractionation, tumor spheroid assay, and Western blot analysis. CTC and YQ contributed to xenografted tumors. YCW, JMS, HJK and DKA participated in the discussion and interpretation of results. GLM and AC provided key reagents. LG and LM contributed to conceptualization. YCW and DKA summarized all the data and wrote the manuscript with input from all authors.

## MATERIALS AND METHODS

### Cell lines, constructs, and treatments

Human breast cancer MDA-MB-231, MDA-MB-468, MCF7, BT-549, Hs578T and embryonic kidney HEK-293T cells were purchased from American Type Culture Collection and cultured with Dulbecco’s Modified Eagle’s Medium (DMEM, Cat# 10-013-CV, Corning) containing fetal bovine serum (10%, Cat# 10437028, ThermoFisher) and 1X antibiotic and antimycotic solution (Cat# 30-004-CI, Corning). Arginine-free media for BT-549, HEK-293T, Hs578T, MCF7, MDA-MB-231 and MDA-MB-468 cells were prepared from DMEM for SILAC (Cat# 88364, ThermoFisher) supplemented with dialyzed FBS (10%, Cat# 26400-044, ThermoFisher), 1X antibiotic and antimycotic solution (Cat# 30-004-CI, Corning), and L-lysine (146.2 μg/ml, Cat# L-9037, Millipore Sigma). For arginine removal, cells were washed with warm arginine-free medium once then incubated in arginine-free medium for indicated times. FLAG and FLAG-tagged PCNA expressing lentiviral plasmids were purchased from GeneCopoeia (EX-NEG-Lv203 and EX-B0066-Lv203). The lentiviral constructs harboring siRNA-resistant pHAGE-HLTF-WT and R71E Hiran mutant were (Taglialatela et al., 2017) were gifts from Dr. Alberto Ciccia and Dr. Angelo Taglialatela. The isogenic pair of HEK-293T-PCNA-WT and HEK-293T-PCNA-K164R cells (Thakar et al., 2020) were obtained from Dr. George-Lucian Moldovan. Other chemicals used in this study include L-Azidohomoalanine (Cat# C10102, ThermoFisher), L-arginine (Cat# A8094, Millipore Sigma), thymidine 5’-monophosphate disodium salt hydrate (Cat# T7004, Millipore Sigma), hydroxyurea (Cat# 102023, MP Biomedicals), (+)-Camptothecin (CPT, Cat# AC276721000, ACROS Organics), BAY 1895344 (Cat# S8666, Selleckchem), Rabusertib (Cat# S2626, Selleckchem), siHLTF (Cat# SASI_Hs01_00162286 and SASI_Hs01_00029458, Sigma Aldrich and Cat# s13138, ThermoFisher), siFLASH (Cat# SASI_Hs02_00343647 and SASI_Hs01_00152963, Sigma Aldrich), siSLBP (Cat# SASI_Hs01_00147022 and SASI_Hs01_00147023, Sigma Aldrich), siSHPRH (Cat# SASI_Hs02_00310973 and SASI_Hs02_00310974, Sigma Aldrich), siSMARCAL1 (Cat# SASI_Hs02_00328762 and SASI_Hs02_00328763, Sigma Aldrich), siZRANB3 (Cat# SASI_Hs01_00109802 and SASI_Hs01_00109803, Sigma Aldrich). AOH1996 (*Gu et al., 2019*) was a gift from Drs. Long Gu and Linda Malkas. pMD2.G (1 µg; Addgene, Cat# 12259) and psPAX2 (2 µg; Addgene, Cat# 12260) were co-transfected with lentiviral expression construct (3 µg) using Lipofectamine 2000 (14 µl; Life Technologies, 11668-019) into HEK-293T cells to generate lentivirus, as described previously (Cheng et al., 2018). Medium was replaced at 24 h post transfection, and viral medium was collected at 72 h post transfection. Viral medium was filtrated with 0.45 µm filter (Millipore Sigma, Cat# SLHAM33SS) and concentrated with Ultra-15 centrifugal filter device (Millipore Sigma, Cat# UFC903024). Concentrated lentivirus-containing medium was recovered and used to transform desired cell lines in the presence of polybrene (10 µg/ml, Millipore Sigma, Cat# TR-1003-G). Clonal cell lines were selected by pre-determined concentration of puromycin (0.5, 1 and 2 µg/ml; Sigma, Cat# P8833) for respective cell line and overexpression of desired protein was confirmed by Western blotting.

### Western blotting

Total cell lysates were prepared by lysing cells with sample buffer and analyzed by Western blotting. Following polyacrylamide gel electrophoresis, separated proteins were transferred to polyvinylidene difluoride (PVDF, Millipore) membranes. In general, the PVDF membranes with transferred proteins were blocked with nonfat milk (5%) in PBST (137 mM NaCl, 10 mM phosphate, 2.7 mM KCl, 0.1% Tween-20, pH 7.4) for 40 mins. Proteins of interest were identified with an anti-CD44 (1:2000; Cat# 3570, Cell Signaling), anti-acetyl-histone H4 (Lys5) (1:20000; Cat# 07-327, Millipore), anti-acetyl-histone H4 (Lys12) (1:5000; Cat# 07-595, Millipore), anti-acetyl-histone H4 (Lys16) (1:3000; Cat# 07-329, Millipore), anti-Actin (1:10000; Cat# MAB1501, Millipore), anti-ATF4 (1:500; Cat# 11815, Cell Signaling), anti-Chk1 (1:2000; Cat# 2345, Cell Signaling), anti-cyclin A (1:1000; Cat# sc-751, Santa Cruz), anti-cyclin B1 (1:1000; Cat# ADI-KAm-CC195, Enzo), anti-eIF2α (1:2000; Cat# 9722, Cell Signaling), anti-γH2AX (1:3000; Cat# A300-081, Bethyl), anti-histone H3 (1:20000; Cat# 39763, Active Motif), anti-histone H4 (1:10000 or 1:500 for long exposure; Cat# ab177840, abcam), anti-lamin A/C (1:2000, Cat# 4777, Cell Signaling), anti-BRCA2 (1:3000, Cat# ab27976, abcam), anti-PCNA (1:3000, Cat# sc-7907, Santa Cruz), anti-phospho-eIF2α (1:1000; Cat# 9721S, Cell Signaling), anti-phospho-Chk1 (Ser296) (1:2000, Cat# ab79758, abcam), anti-phospho-Chk1 (Ser345) (1:2000; Cat# 2348, Cell Signaling), anti-phospho-histone H3 (Ser10) (1:1000; Cat# 06-570, Millipore Sigma), anti-RPA32/RPA2 (1:1000, Cat# ab2175, abcam), anti-phospho-RPA32 (Ser33) (1:3000; Cat# A300-246A, Bethyl Laboratories), anti-trimethyl-histone H3 (Lys4) (1:2000; Cat# 9751P, Cell Signaling), or anti-trimethyl-histone H3 (Lys9) (1:2000; Cat# 61013, Active Motif), anti-ubiquityl-PCNA (Lys164) (1:1000; Cat# 13439, Cell Signaling), anti-ubiquitin Lys63-specific (1:1000; Cat# 5621S, Cell Signaling), anti-SMARCA3 (HLTF, 1:3000, Cat# A300-230A, Bethyl laboratories), anti-Rad6 (1:3000; Cat# ab31917, abcam), anti-Rad18 (1:3000; Cat# ab188235, abcam), anti-RFC5 (1:1000; Cat# 10385, Proteintech), anti-RFC2 (1:1000; Cat# 10410, Proteintech), anti-BRD4 (1:1000; Cat# 28486, Proteintech), anti-ATAD5 (1:1000; Cat# ab72111, abcam), anti-flag (1:5000; Cat# 66008-3-Ig, Proteintech) antibody, anti-FLASH (1:200; Cat# 43339, Cell Signaling), anti-SLBP (1:1000; Cat# HPA019254, Millipore Sigma), anti-SHPRH (1:1000; Cat# sc-514395, Santa Cruz), anti-SMARCAL1 (1:1000; Cat# sc-376377, Santa Cruz), anti-ZRANB3 (1:1000; Cat# NBP2-93301, Novus Biologicals), anti-phospho-Akt (Ser473) (1:500; Cat# 9271, Cell Signaling), anti-Akt (1:1000; Cat# 9272, Cell Signaling), respectively. Following incubation with primary antibodies (in PBST with non-fat milk (5%)) at 4 °C overnight, PVDF membranes were washed with PBST 3 times, and then incubated with horse radish peroxidase-linked anti-mouse or anti-rabbit secondary immunoglobulin antibodies (1:10000; Cat# 7076S, Cat# 7074S Cell signaling) in PBST for 40 mins, Blots were developed with homemade ECL (2.5 mM luminol, 044 mM p-coumaric acid, 0.1 M Tris-HCl pH 8.5, 0.1% hydrogen peroxide in 100 ml distilled water). ChemiDoc Touch imaging system (Bio-Rad) was used to visualize signals. The captured images were analyzed with Image Lab Software (Bio-Rad, version 5.2.1).

### Subcellular fractionation

Equal number (3×10^6^) of cells were harvested in ice-cold nuclear extraction buffer (NEB) (400 μl, 10 mM HEPES pH 7.9, 10 mM KCl, 1.5 mM MgCl_2_, 0.34 M sucrose, 10% glycerol, 1 mM dithiothreitol, 0.1% Triton X-100) containing protease/phosphatase cocktail inhibitors (Thermo Scientific, Cat# 78446) and incubated on ice for 5 min. 300 μl of whole cell lysates were centrifuged at 1300 x g for 4 min at 4°C. Supernatants were collected as cytoplasmic fractions and nuclei pellets were washed with NEB twice and re-suspended in 300 μl of NEB. Total cell lysates (remaining 100 μl of whole cell lysates), cytoplasmic and nuclear fractions were mixed with 4x Laemli buffer (v/v) and analyzed with desired antibodies by Western blotting.

### Cell cycle analyses (Flow cytometry)

For double thymidine block, cells were treated with thymidine (2.5 mM; Cat# 194754, MP Biomedicals) for 16 h and washed twice with PBS. Released cells were incubated in complete medium for 8 h and blocked with the 2^nd^-thymidine treatment. Following 2^nd^-thymidine block (16 h), cells were washed twice with PBS and released in regular medium for analyses. To enrich mitotic cells, cells were treated with thymidine (2.5 mM, 24 h) and released in regular medium. At 3 h post-releasing in regular medium, cells were treated with nocodazole (100 ng/ml, Cat# M1404, Millipore Sigma) for 12 h and released in regular medium for analysis. For BrdU pulse-labeling, cells were pulse-labeled with BrdU (10 μM, Cat# B9285, Millipore Sigma, 30 min) before or after treatment under indicated conditions. Cells were collected and fixed in ethanol (70%) at −20°C overnight. To analyze BrdU incorporated DNA, cells were washed with PBS to remove ethanol and denatured with HCl (2.5 M, 20 min). Cells were washed twice with PBS then stained with FITC conjugated anti-BrdU antibody (BioLegend, Cat# 364104) (1:100) at 37°C for 1 h. After incubation, cells were washed 3 times with PBS and DNA content were visualized by DAPI (1 μM; Cat# D9542, Millipore Sigma, I h) staining at room temperature. Samples were monitored by Attune NxT flow cytometer (Thermofisher) and results were analyzed and quantified by FCS Express (De Novo software). At least 50000 events were collected and 3 independent experiments were performed and analyzed.

### DNA combing assay

Equal number (5×10^4^) of MDA-MB-231 or BT-549 cells were seeded in 60-mm dishes and incubated overnight. Cells were pulse-labeled with CldU (50 μM; Cat# 105478, MP Biomedicals), IdU (250 μM; Cat# 54-42-2, ACROS Organics) and treated as indicated. Cells were washed with PBS and collected. DNA plugs were generated using FiberPrep (Cat# EXT-001, Genomic Vision) and DNA combing was performed using Combicoverslips (Cat# COV-002-RUO, Genomic Vision), according to the manufacturer instructions. DNA combed coverslips were dehydrated with 70%, 90%, and 100% ethanol for 1 min each. DNA combed coverslips were denatured by 0.5 M NaOH plus 1 M NaCl solution for 8 min at room temperature then washed with PBS. The coverslips were dehydrated again with 70%, 90%, and 100% ethanol for 1 min each and air-dried at room temperature for 10 min. Following blocking with 5% BSA in PBS for 5 mins at 37 °C in a humidity chamber, coverslips were incubated with primary antibodies (rat anti-BrdU (BU1/75) (1:500; recognizing CldU; Cat# NB500-169, Novus) and mouse anti-BrdU (B44) (1:500; recognizing ldU; Cat# 347580, BD) with 5% BSA in PBS) for 2 h at 37°C in a humidity chamber. Following incubation, coverslips were washed 3 times with PBST (0.05% Tween 20) then dehydrated with 70%, 90%, and 100% ethanol stepwise. Coverslips were incubated in PBS containing both goat anti-rat 647 secondary antibody (1:500; Cat# A-21247, ThermoFisher) and goat anti-mouse 488 secondary antibody (1:500; Cat# A-11001, ThermoFisher) for 1 h at 37°C in a humidity chamber. After incubation, coverslips were washed 3 times with PBST (0.05% Tween 20) then dehydrated with 70%, 90%, and 100% ethanol stepwise. Coverslips were mounted with prolong diamond antifade mountant (Cat# P36970, ThermoFisher), and sealed with nail polish. DNA replication tracts were visualized with a fluorescence microscope (Observer II, Zeiss) and ImageJ was used for measuring the length of individual labeled DNA tract.

### Analysis of newly synthesized proteins with AHA labeling

Double thymidine synchronized MDA-MB-231 cells were released in L-Azidohomoalanine (AHA, 200 μM, Cat# C10102, ThermoFisher) containing full or arginine-free medium and cells were harvest in indicated time. Cells were collected and washed with PBS for 3 times. AHA-labeled newly synthesized proteins were analyzed by electrophoresis and detected with anti-biotin antibody (Cat# ab201341, abcam). For visualizing total proteins, whole cell lysates were analyzed by electrophoresis and gels were stained with Coomassie blue and imaged on ChemiDoc Touch imaging system (Bio-Rad). The captured images were analyzed with Image Lab Software (Bio-Rad, version 5.2.1).

To assess levels of specific AHA-labeled proteins, MDA-MB-231 cells were labeled with AHA as described above and lysed in RIPA buffer (1% sodium deoxycholate, 50 mM HEPES, 150 mM NaCl, 1% NP-40, 0.1% SDS, 2.5 mM MgCl_2_, 10 mM sodium glycerophosphate, 10 mM sodium biphosphate pH 7.2) containing protease and phosphatase inhibitors (Thermo Scientific, Cat# 78446). Approximately 1.5 mg of proteins from each sample was reduced with TCEP (5 mM, room temperature for 10 min, Cat# 580561, Millipore Sigma) then alkylated with chloroacetamide (20 mM, room temperature for 15 min, Cat# C8625G, TCI America). Alkylated proteins were precipitated with methanol/chloroform/H_2_O and washed 2 times with methanol to remove residual AHA. The precipitated proteins were suspended in suspension buffer (2% SDS, 50 mM HEPES, 150 mM NaCl, pH 7.2, 2.5 mM TCEP) and sonicated briefly with a probe sonicator. Click chemistry reaction was performed with biotin-PEG4-alkyne (100 μM, Cat# 764213, Millipore Sigma), CuSO4 (1 mM, Cat# 422870050, ACROS organics), and L-ascorbate (1 mM, Cat# 352680050, ACROS organics), THPTA ligand (100 μM, Cat# 762342, Millipore Sigma) and mixed on a rotor at room temperature for 2 h. Biotin-alkyne-reacted proteins were precipitated with methanol/chloroform/H_2_O and washed 2 times with methanol to remove residual biotin-alkyne. The precipitated proteins were suspended in suspension buffer (200 μl, 2% SDS, 50 mM HEPES, 150 mM NaCl, pH 7.2, 2.5 mM TCEP) and sonicated briefly with a probe sonicator and diluted to 600 μl with RIPA buffer. High-capacity streptavidin agarose (20 μl, Cat# 20359, ThermoFisher Scientific) was added to each sample and mixed on a rotor at room temperature overnight. The beads were spun down and washed sequentially twice with RIPA buffer (1 ml), twice with 1M KCl, 0.1 M Na_2_CO_3_, 2 M urea in 50 mM HEPES buffer pH 7.5 (1 ml), twice with RIPA buffer (1 ml) and twice with ddH_2_O (1 ml). After washing, samples were mixed with 2x sample buffer and boiled at 100°C for 15 min to extract the proteins. Elution was analyzed by Western blotting with indicated antibodies.

### Protein Transfection

Human recombinant proteins were introduced into cells with ProteoJuice protein transfection reagent (Cat# 71281-3, MilliporeSigma) according to manufacturer’s instruction. Briefly, recipient cells were seeded in a 6-well plate, washed with PBS for 3 times and maintained in OPTI-MEM. Approximately 5 μg of recombinant human His-tagged histone H4 produced in *E. coli* (Cat# 31493, Active Motif) was mixed with 5 μl of ProteoJuice in 100 μl of OPI-MEM and incubated at room temperature for 20 min. After 20 min of incubation, 900 μl of OPI-MEM was added to make a 1000 μl of mixture. After aspirating the OPTI-MEM from the cells, cells were incubated with the transfection mixture at 37°C (5% CO_2_) for 2-3 h. After incubation, cells were washed 3 times with PBS and incubated in full or arginine-free medium as indicated prior to harvesting or subjected to DNA combing assays.

### Immunoprecipitation

Stable expressing FLAG or FLAG-tagged PCNA HEK293T cells were overexpressed with Myc-tagged ubiquitin plasmid (gift from Dr. Hsiu-Ming Shih) for 24 h prior to the indicated treatments. After treatment, cells were harvested by modified RIPA buffer (50mM Tris-HCl pH7.5, 150 mM NaCl, 5 mM EDTA, 0.1% Triton X-100, and 0.1% NP-40) with protease/phosphatase inhibitor (Cat# 78446, ThermoFisher) and incubated in 4°C on rotating mixer for 15 min. Cell debris were removed by 12000 rpm centrifuge for 10 min in 4°C and supernatants were mixed with anti-DYKDDDDK magnetic agarose (Cat# A36797, ThermoFisher) in 4°C overnight on rotating mixer according to manufacturer’s protocol. Magnetic beads were captured with magnetic rack and washed with modified RIPA buffer for 3 times. Samples were eluted by adding 2X SDS-PAGE Sample Buffer and incubated at 100°C for 15 min. Levels of specific proteins were analyzed by Western blotting. To knockdown HLTF, cells were transfected with siRNA targeting HLTF 24 h prior to Myc-ubiquitin overexpression.

### Accelerated native iPOND (aniPOND)

A rapid and accelerated method modified from isolation of protein on nascent DNA (iPOND) was performed following previous study (Leung et al., 2013). In short, MDA-MB-231 cells were pulse labeled by EdU (10 μM) for indicated time along with indicated treatments. Cells were harvested and nuclear extracts were prepared by using the nuclear extraction buffer as described (Leung et al., 2013). Click reaction was performed to conjugate biotin-azide (Cat# 1265, Click Chemistry Tools) to EdU labeled nascent DNA and sonicated (Bioruptor Pico sonication device). Samples were centrifuged and supernatant was collected and mixed with streptavidin-coated beads (Cat# 20359, ThermoFisher) to pull down biotin-EdU labeled nascent DNA. Beads were centrifuged and washed for 3 times and proteins were eluted by adding 2x sample buffer and boiled at 100°C for 15 min. Western blotting was performed to analyze the proteins on nascent DNA. Cells without Edu labeling (EdU(-)) served as negative control. To compensate for the reduced EdU-labeled track in perturbed cells, the EdU labeling time of -R, HU, and CPT treated cells were extended to 25 min while +R cells were labeled for 15 min (**Fig. 5B**).

### Spheroid formation

Equal number (1.2×10^3^) of cells were seeded in ultralow attachment 96-well plates (Cat# CLS3474, Corning) and incubated for 72 h in a 37°C humidity chamber to form spheroids. Spheroids were washed with medium containing indicated chemical twice then incubated in medium containing same chemical for 2 h at 37°C. After 2 h of incubation, spheroids were washed with complete medium three times and then cultured in complete medium for 8 days. Spheroids were visualized at day 0 and day 8 post-treatment by using Cytation 5 imaging reader (BioTek) and ImageJ was used to process the areas of spheroids.

### Cell viability assay

Equal number (5×10^3^) cells were seeded in 96-well plate and cultured under indicated conditions for different time periods. Acid phosphatase (ACP) assay was performed to measure cell viability. Cells were lysed with ACP1 solution (100 μl, 0.1 M sodium acetate, 0.1% Triton X-100, pH 5.7) containing p-Nitrophenyl Phosphate (12 mM) per well and kept in a 37°C incubator. Following 30 min of incubation, the reaction was stopped with NaOH (10 μl, 1 M) and absorbance at 410 nm was recorded using Synergy H1 hybrid multi-mode reader (BioTek). The value of each group measured at day 0 was set as 1.

### Immunofluorescence staining

To visualize H4K12ac and EdU incorporation in BT-549 spheroids, equal number (8×10^3^) cells were seeded in ultralow attachment 96-well plates for 72 h. Spheroids were incubated in EdU (10 μM)- and Hoechest33342 (10 μM)-containing medium for 1 h, collected, fixed with paraformaldehyde (4%) for 10 min and washed twice with PBS. To generate blocks for cryosection, spheroids were incubated in sucrose (15%) for 30 min, followed by sucrose (30%) for 30 min, and sucrose (30%) containing optimal cutting temperature compound (50%, O.C.T., Tissue-TeK, Cat# 4583) for 30 min. Spheroids were then embedded in O.C.T. (100%) in cryomold and 8 μm thick sections were cryosectioned using a cryostat (Leica, CM3050S). EdU incorporation was stained according to manufacturer’s protocol (EdU kit for imaging, Cat# C10640, ThermoFisher). Following washing with PBS 3 times and blocking with BSA (4%) in 0.2% Tween-20 in PBS (PBST (0.2%)) for 30 min in a 37°C humidity chamber, slides were stained with an anti-H4K12ac antibody (1:500; Cat# 07-595, Millipore Sigma) in PBST with BSA (2%) slides in a 37°C humidity chamber for 2 h. Slides were washed 3 times with PBST (0.2%) then incubated with a goat anti-rabbit 488 secondary antibody (1:500; Cat# A-11034, ThermoFisher) in PBST with BSA (2%) in a 37°C humidity chamber. Slides were washed 3 times with PBST (0.2%) and mounted with prolong diamond antifade mountant (Cat# P36970, ThermoFisher), and then sealed with nail polish.

To assess H4K12ac signals in BT-549 spheroids, equal number (8×10^3^) cells were seeded in ultralow attachment 96-well plates and incubated with complete medium in a 37°C humidity chamber for 72 h to form spheroids. Cryosectioning and immunofluorescence staining were performed as described above. Briefly, cryosections were probed with an anti-H4K12ac antibody (1:500; Cat# 07-595, Millipore Sigma) and followed with incubation with a goat anti-rabbit 488 secondary antibody (1:500; Cat# A-11034, ThermoFisher). H4K12ac-positive cells were visualized using a fluorescence microscope (Observer II, Zeiss; LSM 700 Confocal microscope, Zeiss). ImageJ was used for processing the images in each cryosection. For visualizing histone H4K12ac signals in Hs578T or MCF7 spheroids, 3×10^4^ cells were seeded in ultralow attachment 96-well plates and incubated in complete medium to form spheroids in a 37°C humidity chamber. After 5 days of incubation, tumor spheroids were collected and fixed with paraformaldehyde (4%) for 10 min and washed twice with PBS. Cryosectioning and immunofluorescence staining were performed as described above.

To assess H4K12ac and γH2AX signals in BT-549 spheres, 8×10^3^ cells were seeded in ultralow attachment 96-well plates and incubated in complete medium for 72 h to form spheres. Cryosection and immunofluorescence staining were performed as described above. Briefly, sections were probed with an anti-H4K12ac antibody (1:500; Cat# 07-595, Millipore Sigma) or an anti-H2A.X phosphor (Ser139) antibody conjugated with Alexa Fluor 647 (1:500; Cat# 613408, BioLegend). For H4K12ac visualization, after primary antibody incubation, sections were incubated with a goat anti-rabbit 488 secondary antibody (1:500; Cat# A-11034, ThermoFisher). Samples were imaged by a fluorescence microscope (Observer II, Zeiss; LSM 700 Confocal microscope, Zeiss), and ImageJ was used for evaluating the overall intensity of indicated signals in each sphere.

### Chromatin-bound RPA and γH2AX detection

MDA-MB-231 cells were pulse labeled with EdU (10 μM) for 1 h then treated with full, arginine-free, or medium containing HU (2 mM), CPT (2 μM), ATR inhibitor (100 nM) or Chk1 inhibitor (1 μM) for 4 h. Cells were trypsinized, washed with PBS and extracted with cold 0.2% Triton X-100 in PBS for 7 min on ice. After extraction, cells were washed with PBS and fixed with 4% paraformaldehyde in PBS (Cat# BM-155, Boston BioProducts). Fixed cells were blocked with 0.1% BSA in PBS and RPA signals were stained followed by DAPI staining. Percentage of RPA-positive or **γ**H2AX-positive cells in S-phase (EdU-positive) was analyzed by Attune NxT flow cytometer (Thermofisher) and results were quantified by FCS Express (De Novo software).

### Immunohistochemistry

Animal experiments were approved by the Institutional Animal Care and Use Committee at City of Hope. Xenograft tumors were formalin-fixed, paraffin embedded (FFPE) sections were prepared as described previously (Cheng et al., 2018). For detecting the proteins of interest in xenograft tumors, anti-histone 4 lysine 12 acetylation (H4K12ac) antibody (1:500; Cat# ab177793, abcam), anti-histone 4 lysine 5 acetylation (H4K5ac) antibody (1:4000; Cat# ab232507, abcam), anti-Ki67 antibody (1:200, Cat# ab15580, abcam), and anti-CD31 antibody (1:100, Cat# 77699, Cell signaling), respectively, were used for immunohistochemical staining by the pathology core at City of Hope. IHC stain was performed on Ventana Discovery Ultra (Ventana Medical Systems, Roche Diagnostics, Indianapolis, USA) IHC automated stainer. Briefly, tissue samples were sectioned at a thickness of 5 μm and mounted on positively charged glass slides. The sections were deparaffinized and rehydrated. The endogenous peroxidase activity was blocked before antigen retrieval. The antigens were sequentially detected with primary antibody incubation and heat inactivation was performed to prevent antibody cross-reactivity between the same species. Following each primary antibody, DISCOVERY anti-Rabbit HQ and DISCOVERY anti-HQ-HRP were incubated. The positive signals were visualized with DISCOVERY ChromoMap DAB and DISCOVERY Purple Kit, respectively, counterstained with hematoxylin (Ventana) and then coverslipped. Slides were scanned by bright-field microscopy and 4 fields of peripheral and non-necrotic core regions were photograph in each section. H4K5ac and H4K12ac signal intensities were quantified by ImagePro Premier 9.0.

### RNA extraction and qRT-PCR

Total RNA was extracted from cells using the Quick-RNA MiniPrep (Zymo Research, Cat# R1055). qRT-PCR was performed by as described previously (Cheng et al., 2018). In brief, the RNA concentration and purity were analyzed by Synergy H1 hybrid multi-mode reader (BioTek). Complementary DNA (cDNA) was synthesized with 1 µg of total RNA by using the iScript cDNA Synthesis Kit (Bio-Rad, Cat# 1708891). cDNA was diluted 20 times and real-time quantitative PCR was performed by using iTaq Universal SYBR Green Supermix (BioRad, Cat# 1725122) with CFX-96 Real-Time PCR Detection System (BioRad). The primers used in this study targeting histone *H4C5* are, forward: 5’-GGTGTCAAGCGCATTTCTGGTC-3’, and reverse: 5’-CGTAGACCACATCCATCGCTGT-3’. qRT-PCR data from 3 biological replicates were calculated using 2^-ΔΔCt^ method (Nature Protocols 2008, 3, 1101-1108) and normalized to 18S RNA. The mean mRNA abundance in the first time point (0 h) was set as 1.

### Statistical analyses

Data are presented as mean and standard error of mean (Mean ± SEM). Statistical analyses were performed using GraphPad Prism 7.0 software (GraphPad Prism Software Inc., San Diego, CA). Normal distribution was confirmed using Shapiro-Wilk normality test before performing statistical analyses. For normally distributed data, comparison between two means were assessed by unpaired two-tailed Student’s *t* test and that between three or more groups were evaluated using one-way analysis of variance followed by Tukey’s post hoc test. A *p*-value of <0.05 was considered statistically significant.

## SUPPLEMENTARY FIGURE LEGENDS

**Supplementary Figure S1.**
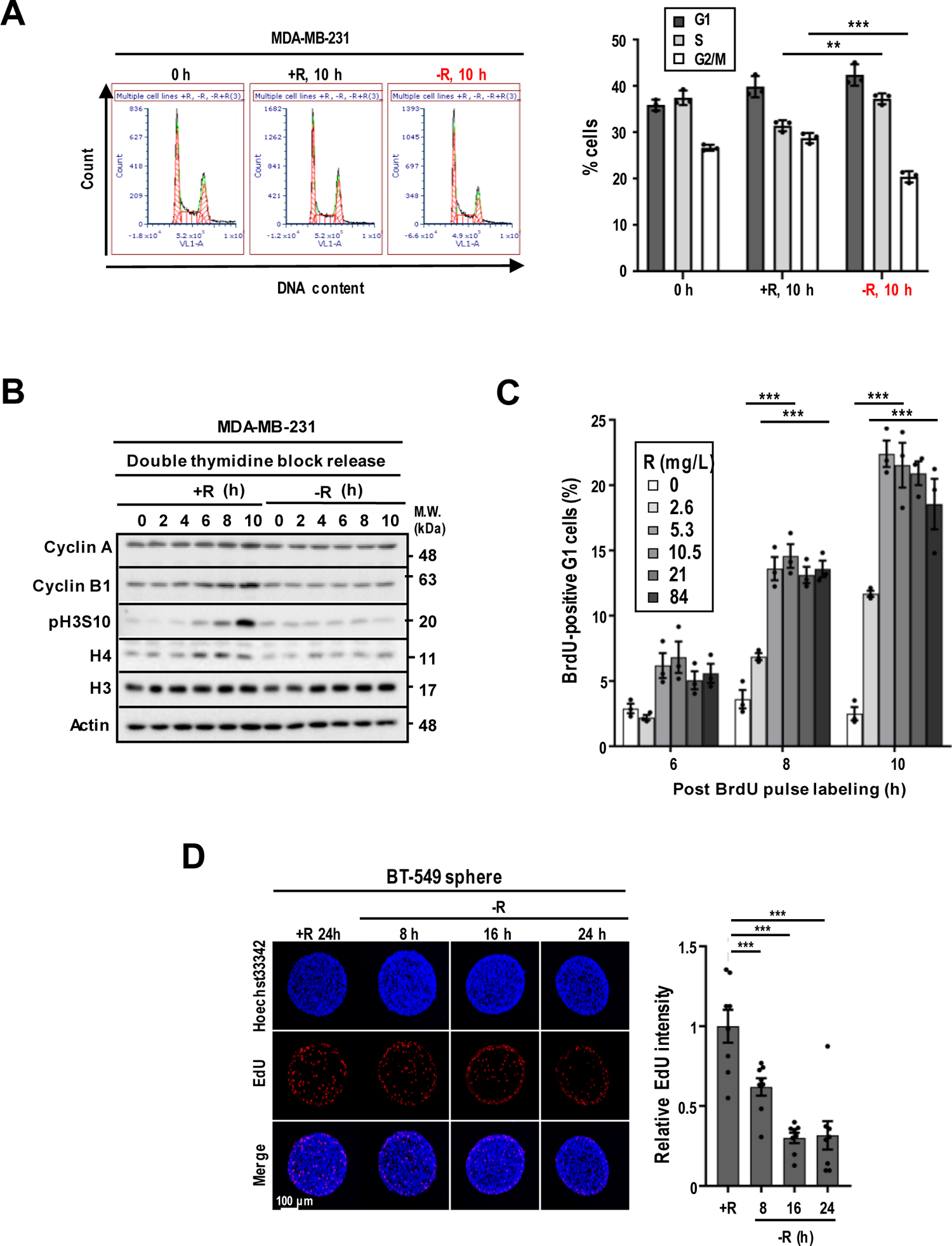
Arginine shortage inhibits S-phase progression. (**A**) Arginine availability affects cell cycle. Flow cytometry analysis of cell cycle phases of MDA-MB-231 cells maintained in medium with or without arginine for 10 h. One representative histogram is shown (N = 3). (**B**) Arginine shortage extends S-phase. Synchronized MDA-MB-231 cells were released in ± R medium and whole cell lysate were collected at indicated time points after release from double thymidine block. Levels of cell cycle and DNA damage markers were analyzed by Western blotting. One representative Western is shown (N = 3). (**C**) Determination on the amount of arginine needed for cell cycle progression. MDA-MB-231 cells were pulse-labeled with BrdU then incubated in medium with an increasing concentration of arginine for indicated times. Percentage of diploid G1 (2N) cells with BrdU-positive signals is shown (N = 3). (**D**) Arginine availability affects DNA replication of tumor spheroids. BT-549 spheroids were incubated with arginine-containing or -free medium for indicated time periods and pulse-labeled with EdU and membrane permeable DNA dye Hoechst 33342 for 1 h. Representative images of DNA (blue), EdU (red) and H4K12ac (green) are shown (*left panel*). Relative EdU intensity quantified and normalized with tumor area is shown (*right panel*; N = 8). (**B, C**) Data are shown as mean ± SEM; two-tailed unpaired Student’s *t*-test; ns: not significant; **: *p*<0.01; ***: *p*<0.005. R: arginine.

**Supplementary Figure S2.**
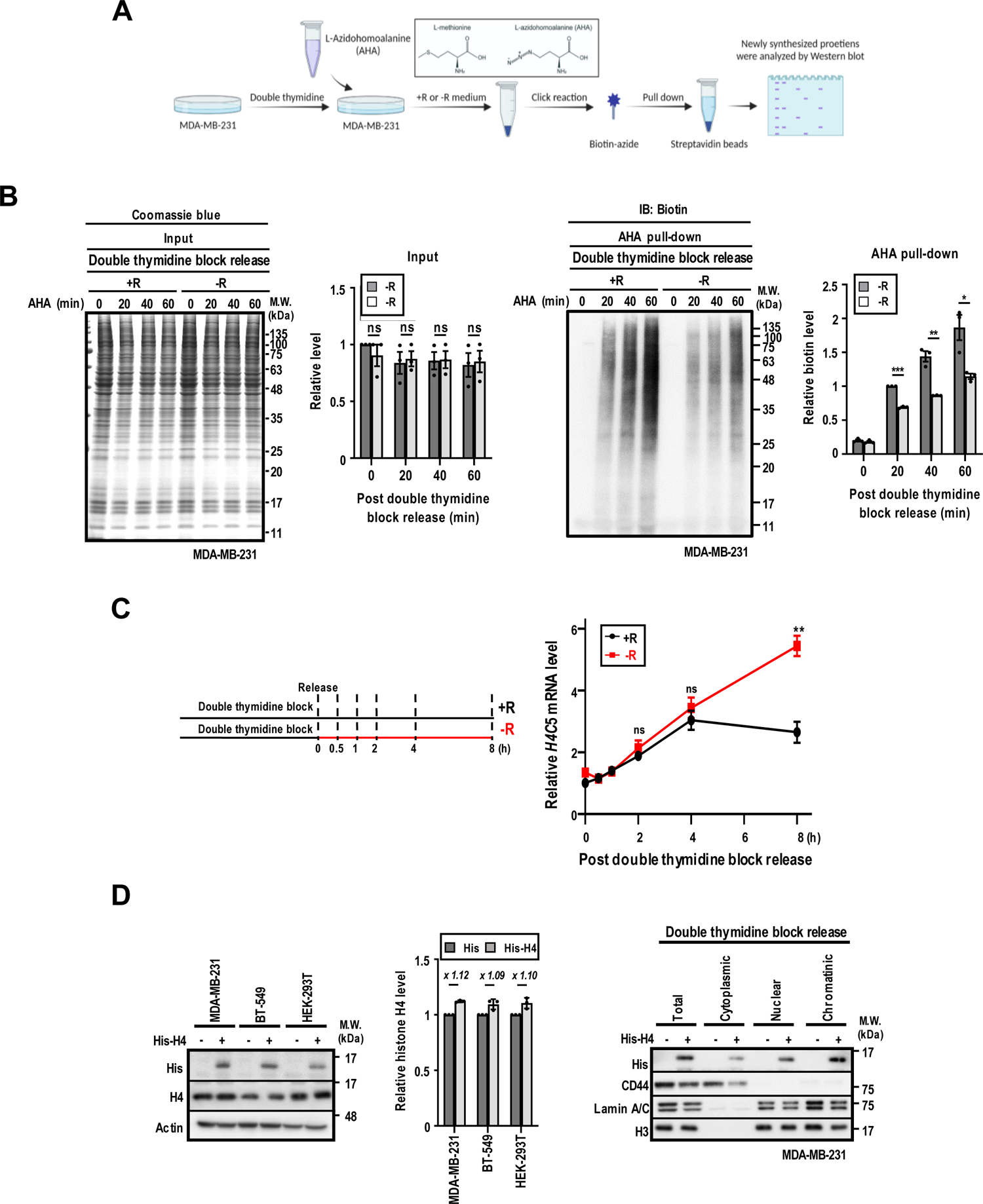
Arginine shortage inhibits histone H4 protein translation. (**A**) Schematic representation of the experimental design for AHA-labeled newly synthesized protein. Created with BioRender.com. (**B**) Arginine is a proteogenic amino acid. MDA-MB-231 cells were synchronized and released in AHA-containing full or arginine-free medium for indicated time. Coomassie blue staining of total proteins is shown (*left panel*). Biotin-conjugated AHA-labeled proteins were detected by anti-biotin antibody and analyzed by SDS-PAGE (*right panel*). (**C**) Arginine shortage does not inhibit histone H4 transcription. MDA-MB-231 cells were synchronized by double thymidine block and released in full or arginine-free medium and cells were collected at indicated time. mRNA levels of histone H4 were analyzed by qRT-PCR. (N = 3) (**D**) Transfected recombinant human His-tagged histone H4 protein is incorporated into chromatin. Recombinant human His-tagged histone H4 protein was transfected in indicated cell lines and Western blot was performed to verify the transfection (*left panel*). Double thymidine block-synchronized MDA-MB-231 cells were transfected with recombinant human His-tagged histone H4 protein. Cells were then harvested, and subcellular fractionations were prepared. CD44 and lamin A/C serve as cytoplasmic and nuclear markers, respectively (*right panel*) (N = 3). R: arginine.

**Supplementary Figure S3.**
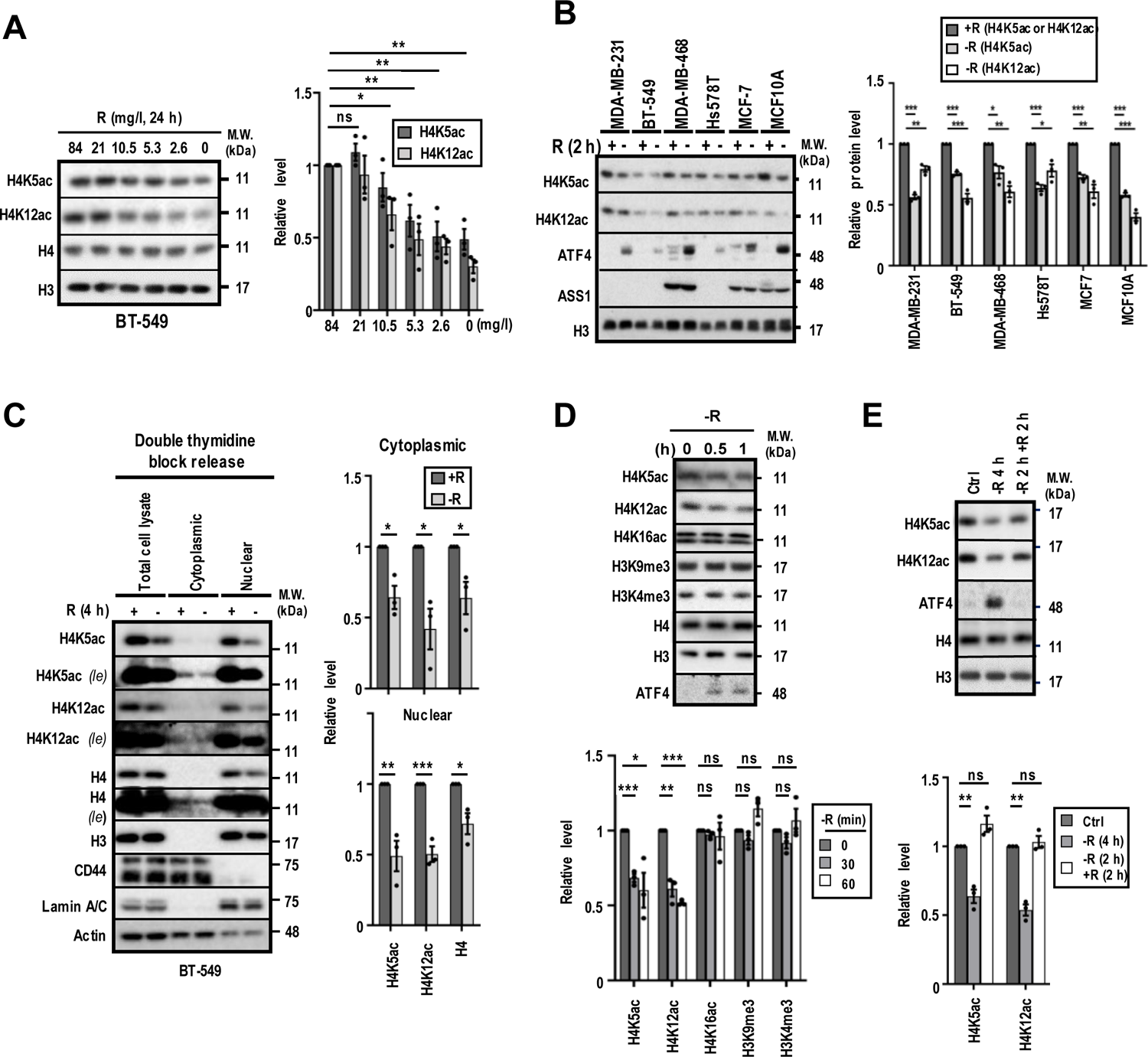
Arginine shortage inhibits newly synthesized histone H4 marks. (**A**) Arginine shortage decreases the steady-state levels of H4K5ac and H4K12ac. BT-549 cells were incubated with medium containing a decreasing concentration of arginine, as indicated, for 24 h. A representative Western is shown (N = 3). (**B**) Arginine shortage decreases H4K5ac and H4K12ac irrespective ASS1 status. ASS1-low breast cancer cell lines (MDA-MB-231, BT-549, Hs578T), ASS1-expressing breast cancer cell lines (MDA-MB-468, MCF7) and ASS1-expressing immortalized mammary epithelial cell line (MCF10A) were incubated with medium ± R for 2 h. Desired protein levels were analyzed by Western blotting. One representative Western (*left panel*) and relative corresponding levels of H4K5ac and H4K12ac signals (*right panel*) are shown. Levels of H4K5ac or H4K12ac in cells treated with complete medium was set as 1, and the relative level of H4K5ac or H4K12ac in arginine-starved cells, after normalization, is shown (N = 3). (**C**) Arginine shortage decreases newly synthesized histone H4. BT-549 cells were synchronized by double thymidine block and released in medium with or without arginine for 4 h. Cells were collected, and subcellular fractions were prepared. CD44 and lamin A/C serve as cytoplasmic and nuclear markers, respectively. One representative Western is shown (N = 3). (**D**) Removal of arginine does not affect H4K16ac, H3K9me3, and H3K4me3 in MDA-MB-231 cells. One representative Western is shown (N=3). (**E**) Arginine shortage regulates the steady-state levels of H4K5ac and H4K12ac in a reversible manner. One representative Western is shown (N = 3). (**A**-**E**) The relative level is calculated as described in **Fig. 2**. Data are shown as mean ± SEM; two-tailed unpaired Student’s *t*-test; ns: not significant; *: *p*<0.05; **: *p*<0.01; ***: *p*<0.005; *le*: long exposure; R: arginine.

**Supplementary Figure S4.**
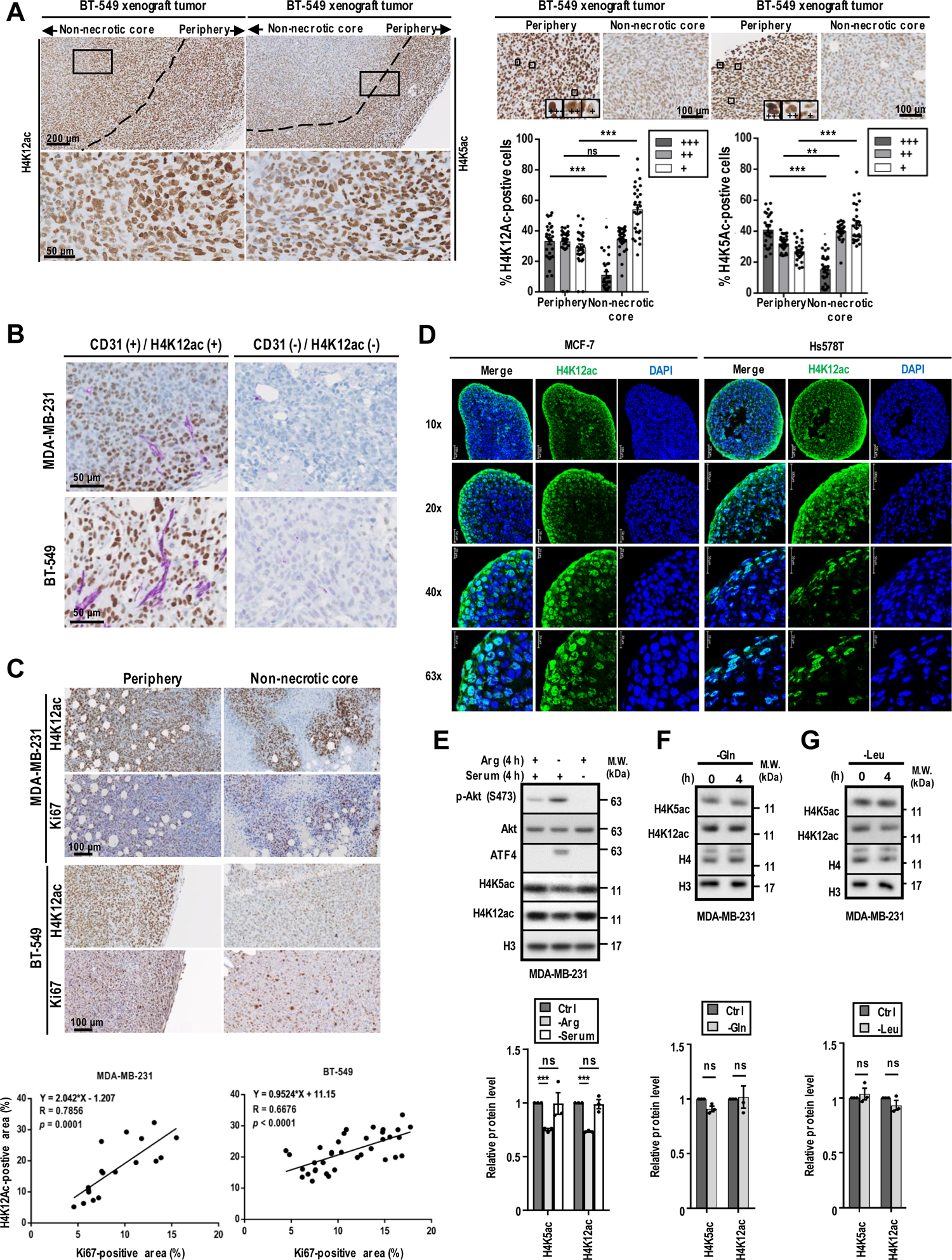
H4K12ac and H4K5ac are gradually decreased inwards to tumor core regions. (**A**) Representative IHC images showing the levels of H4K12ac and H4K5ac decreasing from the peripheral to the core region of BT-549 xenograft tumors. An enlarged view of indicated area (black line, *top left panel*) is shown in *2^nd^ panel*. Black dashed line denotes periphery and non-necrotic core regions. The intensities of H4K12ac- and H4K5ac-positive cells in each IHC images (N = 28) were designated as strong (+++), medium (++), and weak positive (+) (*inlets*), and the respective percentage of cells with indicated H4K5ac or H4K12ac intensity is shown (*right panels*). (**B**) Association of CD31 and H4K12ac in xenograft tumor. Representative images showing adjacent localization of CD31 (purple) and H4K12ac in MDA-MB-231 and BT-549 xenograft tumors visualized by immunohistochemistry. (**C**) Association of H4K12ac and Ki67. Immunohistochemical signals of H4K12ac and Ki67 in MDA-MB-231 and BT-549 (*upper panels*) xenograft tumors were quantified by ImagePro Premier 9.0. A correlation between Ki67 and H4K12ac in MDA-MB-231 (*lower panel*; N = 18, r = 0.7856, 95% CI = 0.5033-0.9164, *p*=0.0001) and BT-549 (*lower panel*; N = 36, r = 0.6676, 95% CI = 0.4343-0.8169, *p*<0.0001) xenograft tumors is shown. (**D**) Localization of H4K12ac (green) in Hs578T and MCF-7 tumor spheroids. Spheroids were collected and visualized by cryosection staining and immunofluorescence microscopy. (**E-G**) Serum, glutamine, and leucine shortage has no effect on H4K5ac and H4K12ac levels. Analysis of H4K5ac and H4K12ac in MDA-MB-231 cells grown under arginine, serum, glutamine, or leucine removal for 4 h. *Left:* Representative Western blot. *Right:* Densitometric quantification of relative levels of H4K5ac and H4K12ac. N = 3 independent experiments. Data are shown as mean ± SEM; two-tailed unpaired Student’s *t*-test; ns: not significant; **: *p*<0.01; ***: *p*<0.005.

**Supplementary Figure S5.**
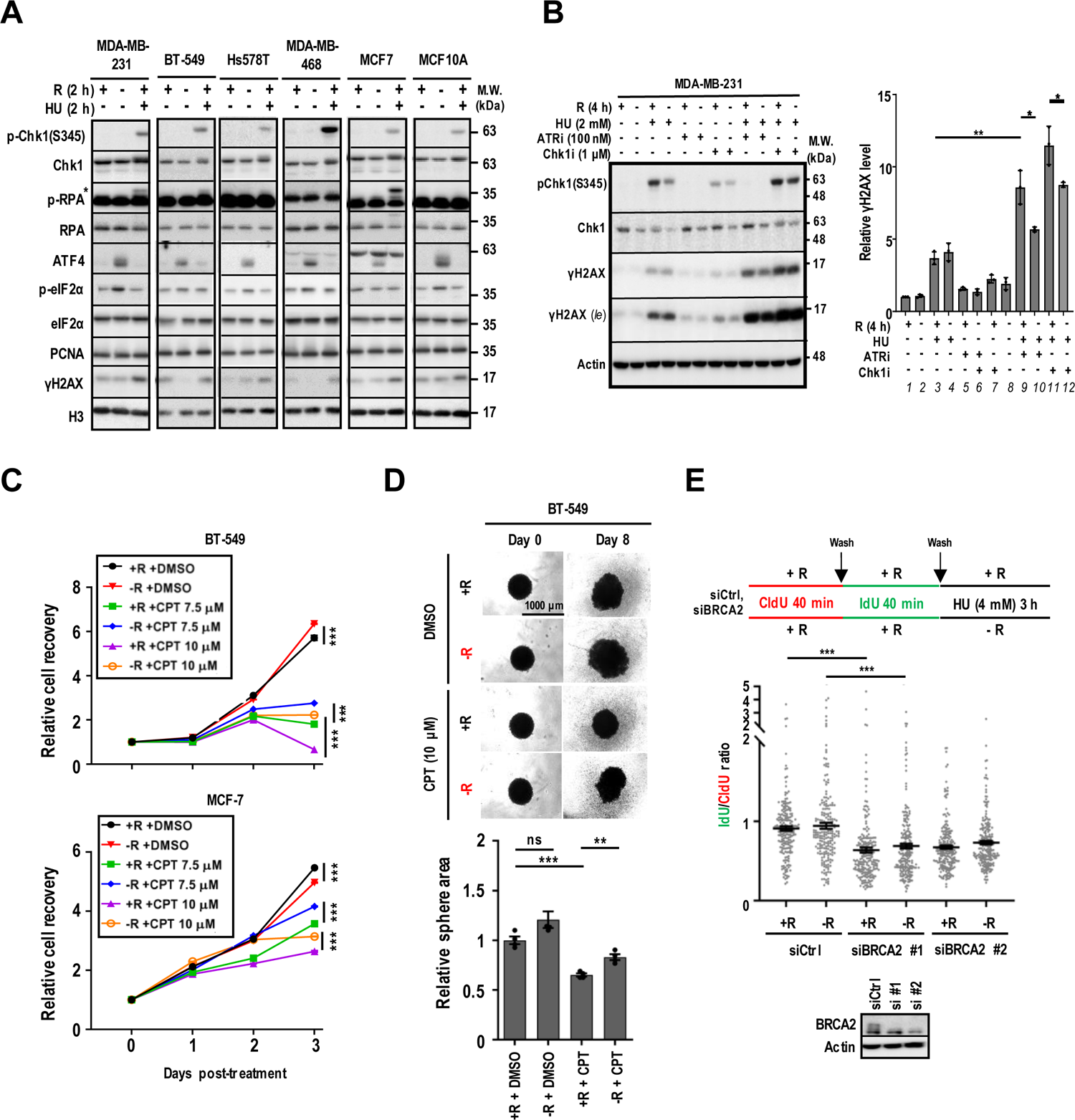
Arginine shortage does not cause canonical replication stress. (**A**) Arginine removal (2 h) does not induce RPA (*upper band, indicated by \**) and Chk1(S345) phosphorylation. A representative Western blot in multiple cell lines is shown (N = 3). (**B**) Arginine shortage (4 h) decreases the DNA damage signaling. One representative Western is shown (*upper panel*). The relative γH2AX (*lower panel*) levels, calculated by dividing densitometric tracing values of γH2AX with that of actin in cells with indicated treatments by the value in cells treated with complete medium (set as 1), after normalizing, are shown (N = 3). (C) Arginine shortage promotes the proliferation of BT-549 and MCF-7 recovered from CPT-treatment. Relative cell recovery of BT-549 and MCF-7 cells after treatment with DMSO, CPT under ± R for 4 h are shown (N = 3) (D) Arginine shortage promotes the proliferation of BT-549 spheroids recovered from CPT-treatment. BT-549 spheroids were treated with vehicle or CPT (10 μM) in ± R for 2 h and recovered in complete medium. Area of tumor spheroids were measured at day 0 and day 8 post-treatment, and relative spheroid area is shown (N=4). (E) Arginine is dispensable for BRCA2-mediated fork protection activities. Fork protection assay was performed by using CldU and IdU to pulse-label ongoing fibers and subjecting MDA-MB-231 cells with high concentration of HU (4 mM) with or without arginine for 3 h. Ratio of IdU/CldU is shown (*lower panel*). BRCA2 was knocked down by 2 siRNA (#1 and #2) and was treated as above. At least 200 tracts were analyzed per sample. Data are shown as mean ± SEM; two-tailed unpaired Student’s *t*-test; ns: not significant; *: *p*<0.05; **: *p*<0.01; ***: *p*<0.005; R: arginine.

**Supplementary Figure S6.**
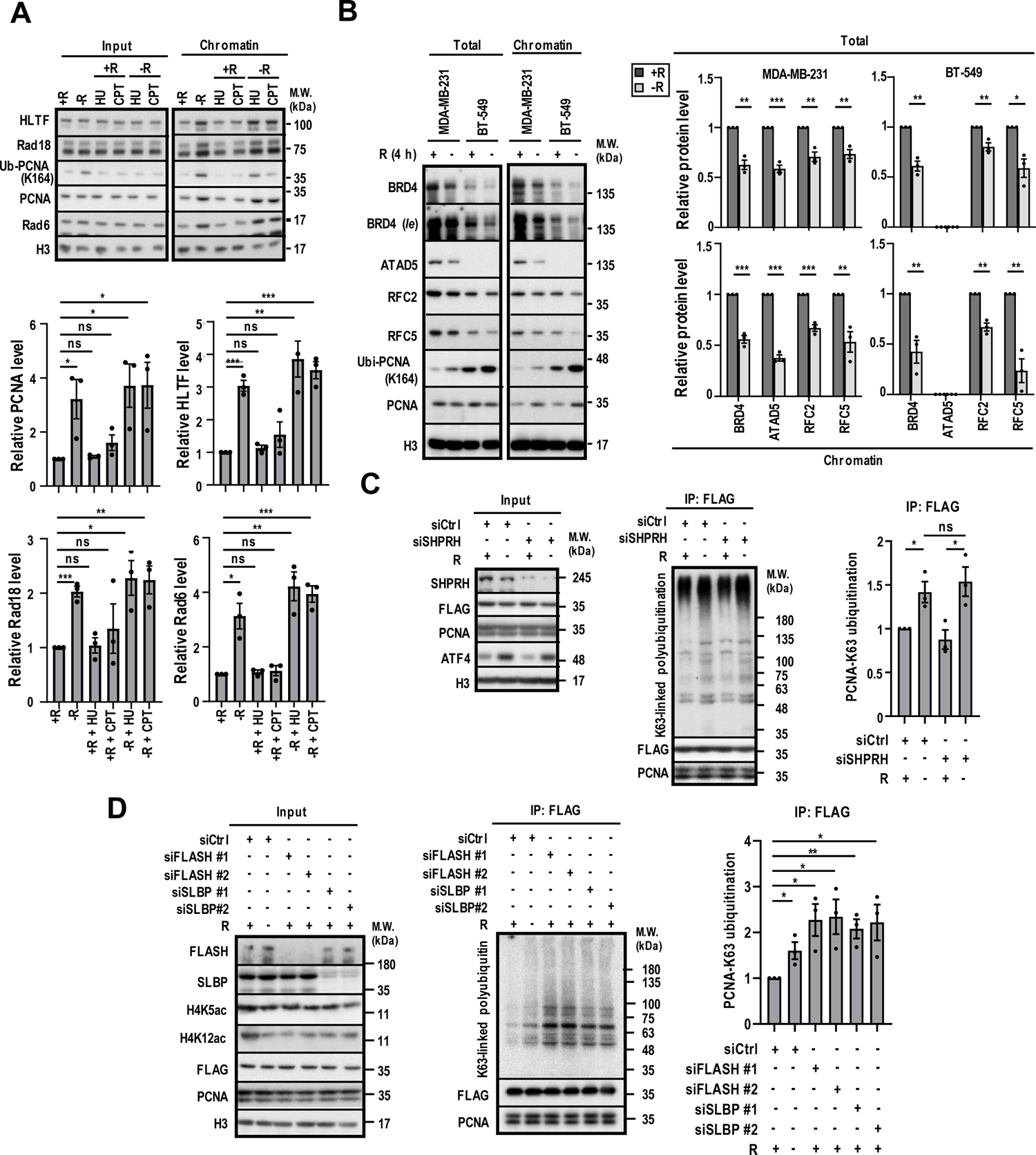
Arginine shortage modulates the abundance of PCNA and its unloading proteins on chromatin. **(A)** Arginine shortage increases the levels of PCNA ubiquitination on chromatin. MDA-MB-231 cells were treated with indicated treatments (HU, 50 µM; CPT, 100 nM) for 4 h and chromatin fractionation was performed to extract chromatin-bound proteins. The levels of indicated proteins were analyzed by Western blotting. A representative Western blot is shown (N = 3). (**B**) Arginine shortage decreases the levels of PCNA unloading-associated proteins. MDA-MB-231 and BT-549 cells were deprived of arginine for 4 h and chromatin fractionation was performed to extract chromatin-bound proteins. The levels of indicated proteins were analyzed by Western blotting. A representative Western blot in multiple cell lines is shown (N = 3). (**C)** Arginine shortage induces PCNA K63-linked polyubiquitination independent of SHPRH. Western blot analysis of immunoprecipitated PCNA-associated proteins from Myc-ubiquitin transfected FLAG-PCNA-overexpressed HEK293T cells and subjected to indicated treatments. Representative Western blots are shown for the indicated antibodies. (N = 3) (**D**) Suppression of histone synthesis induces PCNA K63-linked polyubiquitination. Western blot analysis of immunoprecipitated PCNA-associated proteins from Myc-ubiquitin transfected FLAG-PCNA-overexpressed HEK293T cells and subjected to indicated treatments. Representative Western blots are shown for the indicated antibodies (N = 3). Data are shown as mean ± SEM; two-tailed unpaired Student’s *t*-test; ns: not significant; *: *p*<0.05, **: *p*<0.01; ***: *p*<0.005; *le*: long exposure; R: arginine.

**Supplementary Figure S7.**
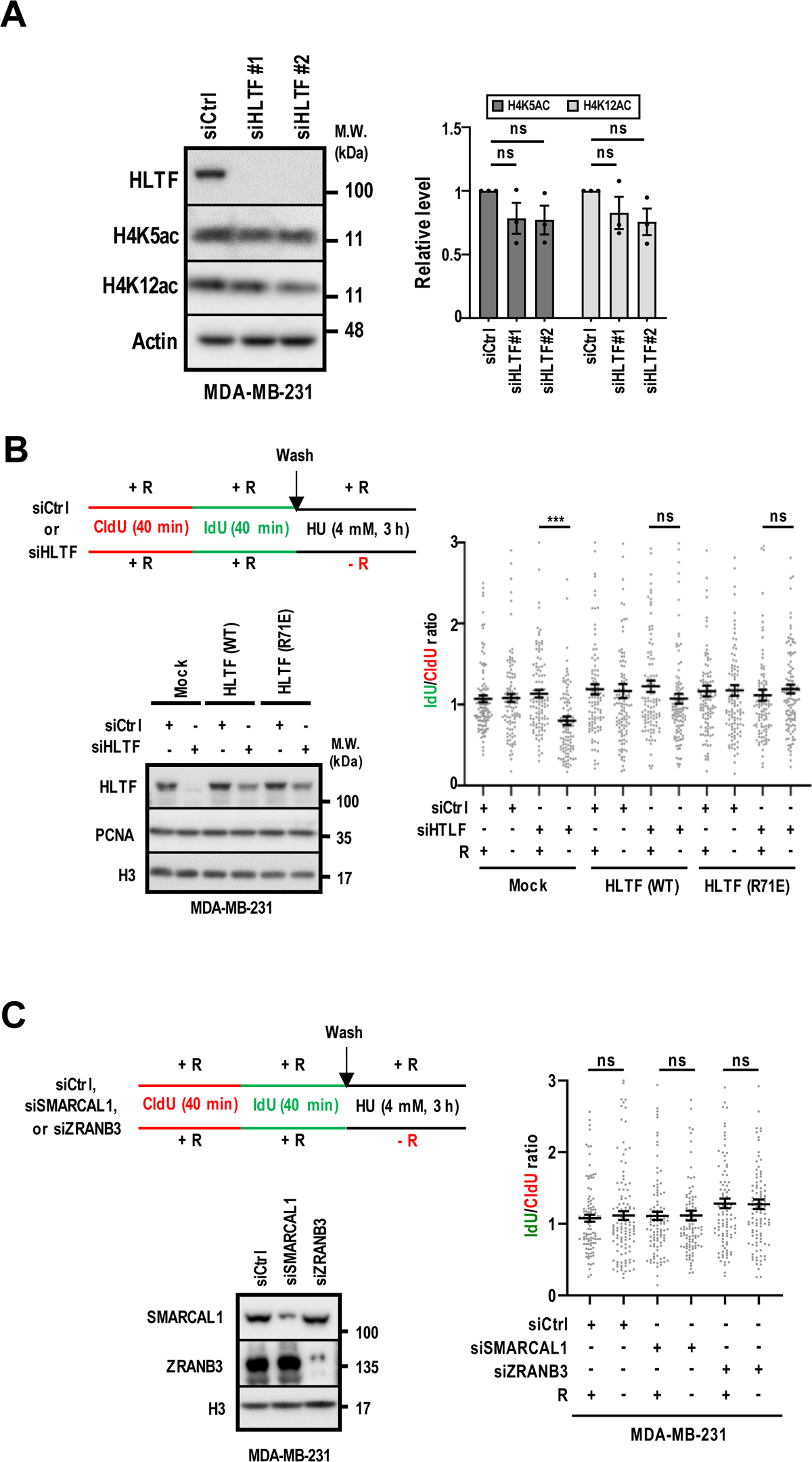
Function of HLTF in arginine-deprived cells. (**A**) Knockdown of HLTF has no effects on H4K5ac and H4K12ac levels. Levels of H4K5ac and H4K12ac in HLTF-knockdown MDA-MB-231 cells were analyzed by Western blot (N = 3). (**B**) HLTF overexpression rescues arginine shortage-induced fork degradation. The effect of wild-type (WT) HLTF and Hiran mutant (R71E) HLTF on replication fork stability under ± arginine upon prolonged HU-treatment was analyzed by DNA combing assay. *Top left:* Experimental outline. *Bottom left:* Representative Western blot analysis of endogenous HLTF and siRNA resistant exogenous HLTF. *Right*: Quantification of IdU/CldU ratio. Each dot represents one fiber. At least 100 tracts were analyzed per sample. (**C**) Knockdown of SMARCAl1 or ZRANB3 has no effect on fork stability in arginine-starved cells. The effect of SMRCAL1- or ZRANB3-knockdown on replication fork stability under ± arginine upon on prolonged HU-treatment was analyzed by DNA combing assay. *Top left:* Experimental outline. *Bottom left:* Representative Western blot analysis of SMARCAL1 and ZRANB3 levels. *Right*: Quantification of IdU/CldU ratio. Each dot represents one fiber. At least 100 tracts were analyzed per sample. Data are shown as mean ± SEM; two-tailed unpaired Student’s *t*-test; ns: not significant; ***: *p*<0.005.

**Supplementary Figure S8.**
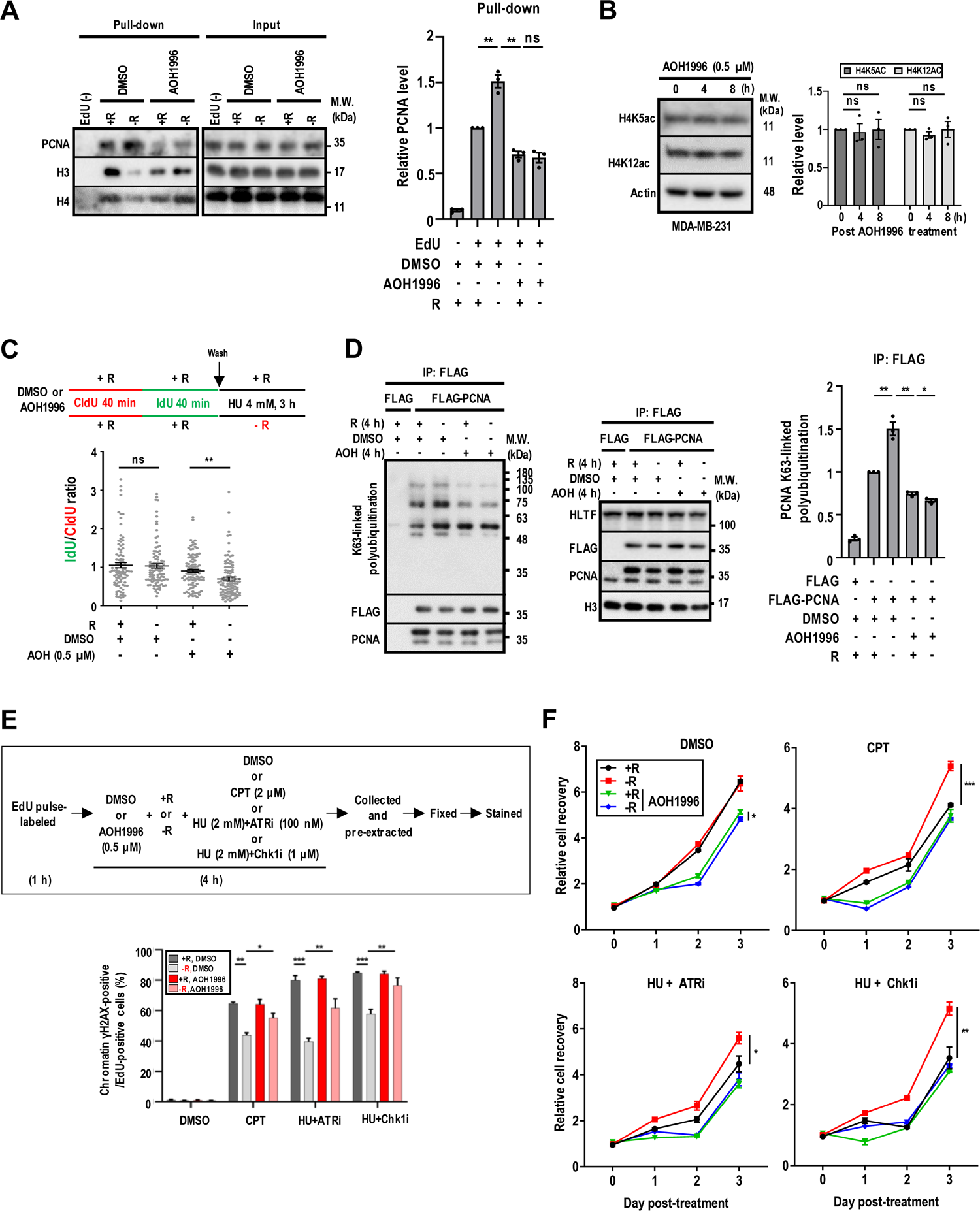
Effects of AOH1996 on arginine-deprived cells. (**A**) AOH1996 treatment prevents PCNA accumulation at the replication fork. The effect of AOH1996 on PCNA deposition on the nascent DNA strands in the presence and absence of arginine was analyzed by aniPOND. *Left:* Representative Western blot. *Right:* Densitometric quantification of relative levels of PCNA in pull-down samples. N = 3 independent experiments. (**B**) AOH1996 has no effect on H4K5ac and H4K12ac levels. Levels of H4K5ac and H4K12ac in MDA-MB-231 cells treated with AOH1996 (0.5 µM) for indicated time were analyzed by Western blotting. *Left:* Representative Western blot. *Right:* Densitometric quantification of relative levels of H4K5ac and H4K12ac. N = 3 independent experiments. (**C**) AOH1996 induces fork degradation in arginine shortage cells. The effect of AOH1996 (0.5 µM) on replication fork stability under ± arginine and exposure to HU was analyzed by DNA combing assay. *Top*: Experimental outline. *Bottom*: Quantification of IdU/CldU ratio. Each dot represents one fiber. At least 100 tracts were analyzed per sample. (**D**) AOH1996 treatment abolishes K63-linked polyubiquitination of PCNA in arginine starved cells. Arginine shortage-dependent K63-linked polyubiquitination of PCNA in control and AOH1996 (0.5 µM)-treated cells (4 h) was analyzed by immunoprecipitation followed by Western blotting. *Left:* Representative Western blot. *Right:* Densitometric quantification of relative levels of K63-linked polyubiquitinated PCNA in pull-down samples. N = 3 independent experiments. (**E**) AOH1996 reverses arginine shortage-attenuated chromatin-bound γH2AX upon genotoxic agents. Vehicle-or AOH1996-treated (0.5 µM) MDA-MB-231 cells were treated with HU (2 mM), HU (2mM) + ATRi (100 nM), HU (2 mM) + Chk1i (1 μM) or CPT (2 μM) under ± arginine for 4 h. Cells were pre-extracted, and chromatin bound γH2AX-positive cells in S-phase were analyzed by flow cytometry (N = 3). (**F**) AOH1996 reverses arginine shortage-promoted cell recovery. Relative cell recovery was analyzed in vehicle- and AOH1996-treated (0.5 µM) MDA-MB-231 cells after treatment with DMSO, CPT, HU + ATRi or HU + Chk1i under ± arginine for 4 h. N = 3 independent experiments. Data are shown as mean ± SEM; two-tailed unpaired Student’s *t*-test; ns: not significant; *: *p*<0.05, **: *p*<0.01; ***: *p*<0.005; CPT: camptothecin; HU: hydroxyurea; R: arginine.

**Supplementary Figure S9.**
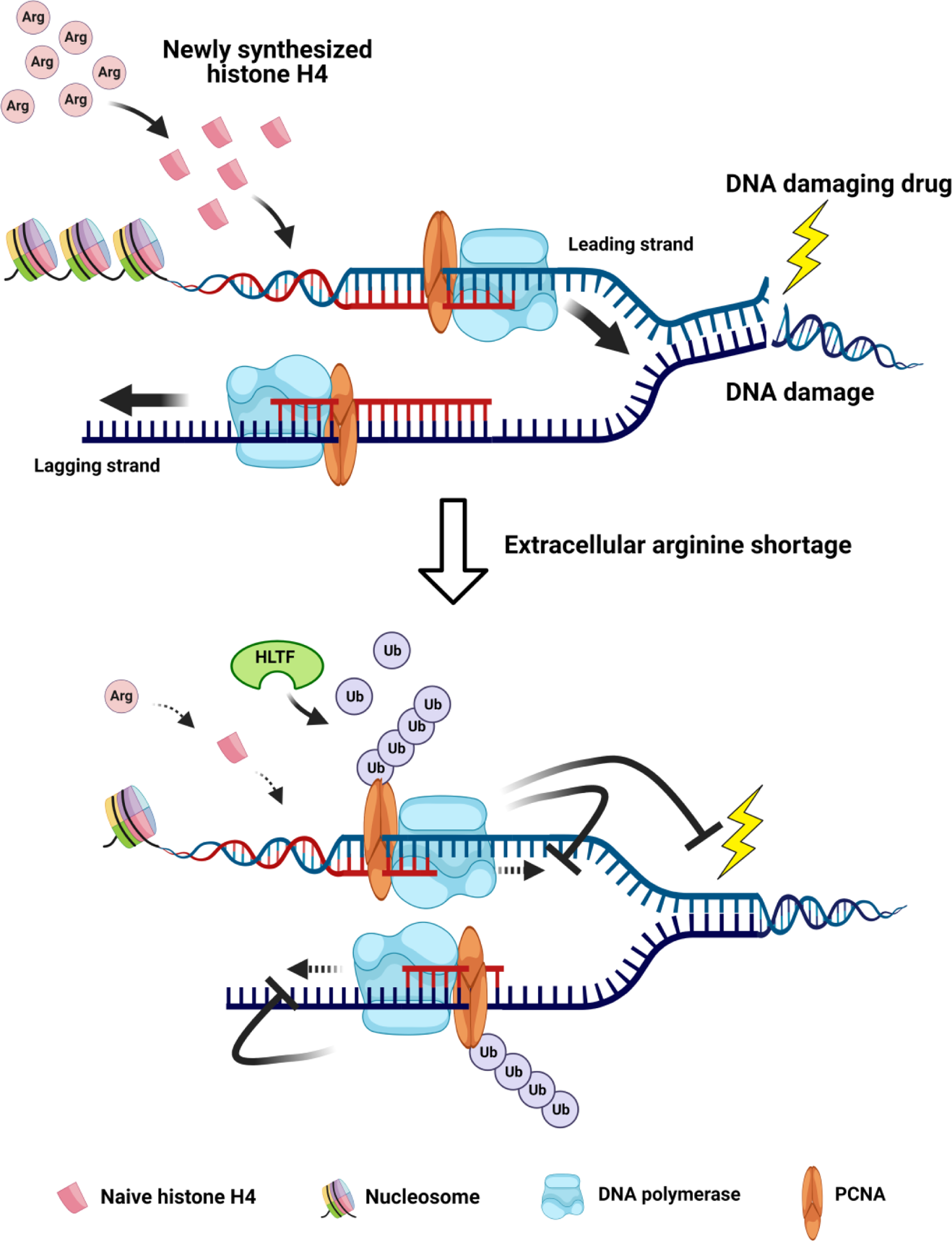
Proposed model for how arginine availability regulates DNA replication and DNA damage. Extracellular arginine shortage limits newly synthesized histone H4 for re-assembly of chromatin at nascent DNA strands, generating instant slowdown of replication forks. In addition, increased PCNA loading and HLTF-mediated K63-linked polyubiquitination for additional constraint elicited by arginine shortage on fork elongation. On the other hand, arginine shortage promotes resistance to genotoxin-induced DNA damage by preventing DNA replication forks from encountering obstacles. Please see discussion for more details. The diagram is created with BioRender.com.

## REFERENCES

Alabert, C., Jasencakova, Z., and Groth, A. (2017). Chromatin Replication and Histone Dynamics. Adv Exp Med Biol 1042, 311–333.

Arbel, M., Choudhary, K., Tfilin, O., and Kupiec, M. (2021). PCNA Loaders and Unloaders-One Ring That Rules Them All. Genes (Basel) 12.

Armstrong, C., and Spencer, S.L. (2021). Replication-dependent histone biosynthesis is coupled to cell-cycle commitment. Proc Natl Acad Sci U S A 118.

Bai, G., Kermi, C., Stoy, H., Schiltz, C.J., Bacal, J., Zaino, A.M., Hadden, M.K., Eichman, B.F., Lopes, M., and Cimprich, K.A. (2020). HLTF Promotes Fork Reversal, Limiting Replication Stress Resistance and Preventing Multiple Mechanisms of Unrestrained DNA Synthesis. Mol Cell 78, 1237–1251 e1237.

Berti, M., Cortez, D., and Lopes, M. (2020). The plasticity of DNA replication forks in response to clinically relevant genotoxic stress. Nature reviews Molecular cell biology 21, 633–651.

Betous, R., Goullet de Rugy, T., Pelegrini, A.L., Queille, S., de Villartay, J.P., and Hoffmann, J.S. (2018). DNA replication stress triggers rapid DNA replication fork breakage by Artemis and XPF. PLoS genetics 14, e1007541.

Blastyak, A., Hajdu, I., Unk, I., and Haracska, L. (2010). Role of double-stranded DNA translocase activity of human HLTF in replication of damaged DNA. Mol Cell Biol 30, 684–693.

Bohnsack, B.L., and Hirschi, K.K. (2004). Nutrient regulation of cell cycle progression. Annual review of nutrition 24, 433–453.

Boroughs, L.K., and DeBerardinis, R.J. (2015). Metabolic pathways promoting cancer cell survival and growth. Nat Cell Biol 17, 351–359.

Bruhl, J., Trautwein, J., Schafer, A., Linne, U., and Bouazoune, K. (2019). The DNA repair protein SHPRH is a nucleosome-stimulated ATPase and a nucleosome-E3 ubiquitin ligase. Epigenetics Chromatin 12, 52.

Chen, C.L., Hsu, S.C., Ann, D.K., Yen, Y., and Kung, H.J. (2021a). Arginine Signaling and Cancer Metabolism. Cancers (Basel) 13.

Chen, C.L., Hsu, S.C., Chung, T.Y., Chu, C.Y., Wang, H.J., Hsiao, P.W., Yeh, S.D., Ann, D.K., Yen, Y., and Kung, H.J. (2021b). Arginine is an epigenetic regulator targeting TEAD4 to modulate OXPHOS in prostate cancer cells. Nature communications 12, 2398.

Cheng, C.T., Qi, Y., Wang, Y.C., Chi, K.K., Chung, Y., Ouyang, C., Chen, Y.R., Oh, M.E., Sheng, X., Tang, Y., et al. (2018). Arginine starvation kills tumor cells through aspartate exhaustion and mitochondrial dysfunction. Commun Biol 1, 178.

Cho, S., Cinghu, S., Yu, J.R., and Park, W.Y. (2011). Helicase-like transcription factor confers radiation resistance in cervical cancer through enhancing the DNA damage repair capacity. J Cancer Res Clin Oncol 137, 629–637.

Choe, K.N., and Moldovan, G.L. (2017). Forging Ahead through Darkness: PCNA, Still the Principal Conductor at the Replication Fork. Mol Cell 65, 380–392.

Cimprich, K.A., and Cortez, D. (2008). ATR: an essential regulator of genome integrity. Nature reviews Molecular cell biology 9, 616–627.

Cuyas, E., Corominas-Faja, B., Joven, J., and Menendez, J.A. (2014). Cell cycle regulation by the nutrient-sensing mammalian target of rapamycin (mTOR) pathway. Methods in molecular biology 1170, 113–144.

Darnell, A.M., Subramaniam, A.R., and O’Shea, E.K. (2018). Translational Control through Differential Ribosome Pausing during Amino Acid Limitation in Mammalian Cells. Molecular cell 71, 229–243 e211.

Denu, J.M. (2019). Histone Acetyltransferase 1 Links Metabolism and Transcription to Cell-Cycle Progression. Molecular cell 75, 664–665.

Forment, J.V., and Jackson, S.P. (2015). A flow cytometry-based method to simplify the analysis and quantification of protein association to chromatin in mammalian cells. Nature protocols 10, 1297–1307.

Gunesdogan, U., Jackle, H., and Herzig, A. (2014). Histone supply regulates S phase timing and cell cycle progression. eLife 3, e02443.

Hammond, C.M., Stromme, C.B., Huang, H., Patel, D.J., and Groth, A. (2017). Histone chaperone networks shaping chromatin function. Nature reviews Molecular cell biology 18, 141–158.

Hoege, C., Pfander, B., Moldovan, G.L., Pyrowolakis, G., and Jentsch, S. (2002). RAD6-dependent DNA repair is linked to modification of PCNA by ubiquitin and SUMO. Nature 419, 135–141.

Hsu, S.C., Chen, C.L., Cheng, M.L., Chu, C.Y., Changou, C.A., Yu, Y.L., Yeh, S.D., Kuo, T.C., Kuo, C.C., Chuu, C.P., et al. (2021). Arginine starvation elicits chromatin leakage and cGAS-STING activation via epigenetic silencing of metabolic and DNA-repair genes. Theranostics 11, 7527–7545.

Kanao, R., and Masutani, C. (2017). Regulation of DNA damage tolerance in mammalian cells by post-translational modifications of PCNA. Mutation research 803-805, 82-88.

Kang, M.S., Kim, J., Ryu, E., Ha, N.Y., Hwang, S., Kim, B.G., Ra, J.S., Kim, Y.J., Hwang, J.M., Myung, K., et al. (2019). PCNA Unloading Is Negatively Regulated by BET Proteins. Cell Rep 29, 4632–4645 e4635.

Kannouche, P.L., Wing, J., and Lehmann, A.R. (2004). Interaction of human DNA polymerase eta with monoubiquitinated PCNA: a possible mechanism for the polymerase switch in response to DNA damage. Molecular cell 14, 491–500.

Kawasumi, M., Bradner, J.E., Tolliday, N., Thibodeau, R., Sloan, H., Brummond, K.M., and Nghiem, P. (2014). Identification of ATR-Chk1 pathway inhibitors that selectively target p53-deficient cells without directly suppressing ATR catalytic activity. Cancer research 74, 7534–7545.

Kelman, Z. (1997). PCNA: structure, functions and interactions. Oncogene 14, 629–640.

Keshet, R., Szlosarek, P., Carracedo, A., and Erez, A. (2018). Rewiring urea cycle metabolism in cancer to support anabolism. Nat Rev Cancer 18, 634–645.

Kile, A.C., Chavez, D.A., Bacal, J., Eldirany, S., Korzhnev, D.M., Bezsonova, I., Eichman, B.F., and Cimprich, K.A. (2015). HLTF’s Ancient HIRAN Domain Binds 3’ DNA Ends to Drive Replication Fork Reversal. Mol Cell 58, 1090–1100.

Kolinjivadi, A.M., Sannino, V., De Antoni, A., Zadorozhny, K., Kilkenny, M., Techer, H., Baldi, G., Shen, R., Ciccia, A., Pellegrini, L., et al. (2017). Smarcal1-Mediated Fork Reversal Triggers Mre11-Dependent Degradation of Nascent DNA in the Absence of Brca2 and Stable Rad51 Nucleofilaments. Mol Cell 67, 867–881 e867.

Komatsu, K., Wharton, W., Hang, H., Wu, C., Singh, S., Lieberman, H.B., Pledger, W.J., and Wang, H.G. (2000). PCNA interacts with hHus1/hRad9 in response to DNA damage and replication inhibition. Oncogene 19, 5291–5297.

Lama-Sherpa, T.D., and Shevde, L.A. (2020). An Emerging Regulatory Role for the Tumor Microenvironment in the DNA Damage Response to Double-Strand Breaks. Mol Cancer Res 18, 185–193.

Lee, J.S., Adler, L., Karathia, H., Carmel, N., Rabinovich, S., Auslander, N., Keshet, R., Stettner, N., Silberman, A., Agemy, L., et al. (2018). Urea Cycle Dysregulation Generates Clinically Relevant Genomic and Biochemical Signatures. Cell 174, 1559–1570 e1522.

Lee, S.W., Zhang, Y., Jung, M., Cruz, N., Alas, B., and Commisso, C. (2019). EGFR-Pak Signaling Selectively Regulates Glutamine Deprivation-Induced Macropinocytosis. Dev Cell 50, 381–392 e385.

Leung-Pineda, V., Ryan, C.E., and Piwnica-Worms, H. (2006). Phosphorylation of Chk1 by ATR is antagonized by a Chk1-regulated protein phosphatase 2A circuit. Molecular and cellular biology 26, 7529–7538.

Leung, K.H., Abou El Hassan, M., and Bremner, R. (2013). A rapid and efficient method to purify proteins at replication forks under native conditions. BioTechniques 55, 204–206.

Liu, G.Y., and Sabatini, D.M. (2020). mTOR at the nexus of nutrition, growth, ageing and disease. Nature reviews Molecular cell biology 21, 183–203.

Liu, K.A., Lashinger, L.M., Rasmussen, A.J., and Hursting, S.D. (2014). Leucine supplementation differentially enhances pancreatic cancer growth in lean and overweight mice. Cancer Metab 2, 6.

MacAlpine, D.M., and Almouzni, G. (2013). Chromatin and DNA replication. Cold Spring Harb Perspect Biol 5, a010207.

Marino-Ramirez, L., Jordan, I.K., and Landsman, D. (2006). Multiple independent evolutionary solutions to core histone gene regulation. Genome biology 7, R122.

Mejlvang, J., Feng, Y., Alabert, C., Neelsen, K.J., Jasencakova, Z., Zhao, X., Lees, M., Sandelin, A., Pasero, P., Lopes, M., et al. (2014). New histone supply regulates replication fork speed and PCNA unloading. J Cell Biol 204, 29–43.

Moldovan, G.L., Pfander, B., and Jentsch, S. (2007). PCNA, the maestro of the replication fork. Cell 129, 665–679.

Nakamura, K., Kustatscher, G., Alabert, C., Hodl, M., Forne, I., Volker-Albert, M., Satpathy, S., Beyer, T.E., Mailand, N., Choudhary, C., et al. (2021). Proteome dynamics at broken replication forks reveal a distinct ATM-directed repair response suppressing DNA double-strand break ubiquitination. Molecular cell 81, 1084–1099 e1086.

Pan, M., Reid, M.A., Lowman, X.H., Kulkarni, R.P., Tran, T.Q., Liu, X., Yang, Y., Hernandez-Davies, J.E., Rosales, K.K., Li, H., et al. (2016). Regional glutamine deficiency in tumours promotes dedifferentiation through inhibition of histone demethylation. Nat Cell Biol 18, 1090–1101.

Papouli, E., Chen, S., Davies, A.A., Huttner, D., Krejci, L., Sung, P., and Ulrich, H.D. (2005). Crosstalk between SUMO and ubiquitin on PCNA is mediated by recruitment of the helicase Srs2p. Molecular cell 19, 123–133.

Peng, M., Cong, K., Panzarino, N.J., Nayak, S., Calvo, J., Deng, B., Zhu, L.J., Morocz, M., Hegedus, L., Haracska, L., et al. (2018). Opposing Roles of FANCJ and HLTF Protect Forks and Restrain Replication during Stress. Cell Rep 24, 3251–3261.

Poillet-Perez, L., Xie, X., Zhan, L., Yang, Y., Sharp, D.W., Hu, Z.S., Su, X., Maganti, A., Jiang, C., Lu, W., et al. (2018). Autophagy maintains tumour growth through circulating arginine. Nature 563, 569–573.

Qiu, F., Chen, Y.R., Liu, X., Chu, C.Y., Shen, L.J., Xu, J., Gaur, S., Forman, H.J., Zhang, H., Zheng, S., et al. (2014). Arginine starvation impairs mitochondrial respiratory function in ASS1-deficient breast cancer cells. Science signaling 7, ra31.

Quinet, A., Tirman, S., Cybulla, E., Meroni, A., and Vindigni, A. (2021). To skip or not to skip: choosing repriming to tolerate DNA damage. Mol Cell 81, 649–658.

Ramaswamy, S., Venkatadri, N., and Bapna, J.S. (1987). An investigation into the possible development of chronic tolerance to analgesia and dependence on prolactin. Fundam Clin Pharmacol 1, 445–449.

Rausch, L.K., Netzer, N.C., Hoegel, J., and Pramsohler, S. (2017). The Linkage between Breast Cancer, Hypoxia, and Adipose Tissue. Frontiers in oncology 7, 211.

Shibahara, K., and Stillman, B. (1999). Replication-dependent marking of DNA by PCNA facilitates CAF-1-coupled inheritance of chromatin. Cell 96, 575–585.

Sirbu, B.M., Couch, F.B., Feigerle, J.T., Bhaskara, S., Hiebert, S.W., and Cortez, D. (2011). Analysis of protein dynamics at active, stalled, and collapsed replication forks. Genes & development 25, 1320–1327.

Smith, S., and Stillman, B. (1989). Purification and characterization of CAF-I, a human cell factor required for chromatin assembly during DNA replication in vitro. Cell 58, 15–25.

Sobel, R.E., Cook, R.G., Perry, C.A., Annunziato, A.T., and Allis, C.D. (1995). Conservation of deposition-related acetylation sites in newly synthesized histones H3 and H4. Proceedings of the National Academy of Sciences of the United States of America 92, 1237–1241.

Taddei, A., Roche, D., Sibarita, J.B., Turner, B.M., and Almouzni, G. (1999). Duplication and maintenance of heterochromatin domains. J Cell Biol 147, 1153–1166.

Taglialatela, A., Alvarez, S., Leuzzi, G., Sannino, V., Ranjha, L., Huang, J.W., Madubata, C., Anand, R., Levy, B., Rabadan, R., et al. (2017). Restoration of Replication Fork Stability in BRCA1- and BRCA2-Deficient Cells by Inactivation of SNF2-Family Fork Remodelers. Mol Cell 68, 414–430 e418.

Thakar, T., Leung, W., Nicolae, C.M., Clements, K.E., Shen, B., Bielinsky, A.K., and Moldovan, G.L. (2020). Ubiquitinated-PCNA protects replication forks from DNA2-mediated degradation by regulating Okazaki fragment maturation and chromatin assembly. Nat Commun 11, 2147.

Ulrich, H.D. (2009). Regulating post-translational modifications of the eukaryotic replication clamp PCNA. DNA repair 8, 461–469.

Ulrich, H.D., and Takahashi, T. (2013). Readers of PCNA modifications. Chromosoma 122, 259–274.

Unk, I., Hajdu, I., Fatyol, K., Hurwitz, J., Yoon, J.H., Prakash, L., Prakash, S., and Haracska, L. (2008). Human HLTF functions as a ubiquitin ligase for proliferating cell nuclear antigen polyubiquitination. Proc Natl Acad Sci U S A 105, 3768–3773.

van Harten, A.M., Buijze, M., van der Mast, R., Rooimans, M.A., Martens-de Kemp, S.R., Bachas, C., Brink, A., Stigter-van Walsum, M., Wolthuis, R.M.F., and Brakenhoff, R.H. (2019). Targeting the cell cycle in head and neck cancer by Chk1 inhibition: a novel concept of bimodal cell death. Oncogenesis 8, 38.

van Toorn, M., Turkyilmaz, Y., Han, S., Zhou, D., Kim, H.S., Salas-Armenteros, I., Kim, M., Akita, M., Wienholz, F., Raams, A., et al. (2022). Active DNA damage eviction by HLTF stimulates nucleotide excision repair. Mol Cell 82, 1343–1358 e1348.

Vujanovic, M., Krietsch, J., Raso, M.C., Terraneo, N., Zellweger, R., Schmid, J.A., Taglialatela, A., Huang, J.W., Holland, C.L., Zwicky, K., et al. (2017). Replication Fork Slowing and Reversal upon DNA Damage Require PCNA Polyubiquitination and ZRANB3 DNA Translocase Activity. Mol Cell 67, 882–890 e885.

Wu, G. (2009). Amino acids: metabolism, functions, and nutrition. Amino Acids 37, 1–17.

Ye, C., Sun, N.X., Ma, Y., Zhao, Q., Zhang, Q., Xu, C., Wang, S.B., Sun, S.H., Wang, F., and Li, W. (2015). MicroRNA-145 contributes to enhancing radiosensitivity of cervical cancer cells. FEBS Lett 589, 702–709.

Zellweger, R., Dalcher, D., Mutreja, K., Berti, M., Schmid, J.A., Herrador, R., Vindigni, A., and Lopes, M. (2015). Rad51-mediated replication fork reversal is a global response to genotoxic treatments in human cells. The Journal of cell biology 208, 563–579.

Zeman, M.K., and Cimprich, K.A. (2014). Causes and consequences of replication stress. Nature cell biology 16, 2–9.

